# Auditory selective attention during a working memory task with melodies: a MEG study

**DOI:** 10.1101/2023.09.19.558422

**Authors:** de la Chapelle Aurélien, Serres-Blain Salomé, ElShafei Alma, Schwartz Denis, Daligault Sébastien, Jung Julien, Ruby Perrine, Caclin Anne, Bidet-Caulet Aurélie

**Affiliations:** Université Claude Bernard Lyon 1, INSERM, CNRS, Centre de Recherche en Neurosciences de Lyon CRNL U1028 UMR5292, F-69500, Bron, France; Donders Institute for Brain, Cognition, & Behavior, Radboud University, Nijmegen, The Netherlands, 6525 EN; CERMEP - Imagerie du Vivant, MEG Department, Lyon, France; Department of Functional Neurology and Epileptology, Hospices Civils de Lyon, Lyon, France; Inserm, INS, Inst Neurosci Syst, Marseille, France; Aix-Marseille Univ, Marseille, France

**Keywords:** short-term memory, attentional filtering, auditory sequences, ERF, theta oscillations, alpha oscillations

## Abstract

Working memory and attention are jointly needed in most everyday life tasks and activities. They have however mostly been studied separately. Here we investigate how auditory working memory and selective attention interact using a recently introduced paradigm (MEMAT) that combines a classic working memory paradigm, the delayed-matching-to-sample task, and selective attention, with distractors presented during the encoding phase. All stimuli are four-tone melodies. Twenty-two participants performed the MEMAT task during MEG recordings. We manipulate the difficulty of the memory task and of attentional filtering. *When memory task difficulty increases*, the amplitude of the CNV in anticipation of the melody to encode increases and the decrease in alpha power during encoding and maintenance in a left fronto-temporal network is reduced. *When attentional filtering difficulty increases*, the amplitude of the sustained evoked response during encoding increases, whereas the differential processing of relevant and irrelevant sounds in auditory areas is less pronounced, and frontal theta power during encoding and maintenance is higher. In the left auditory cortex, we could directly observe the result of the interaction between auditory memory and attention: the facilitation of relevant sound processing in the easy filtering condition was reduced when the memory task difficulty increased. This pattern mirrors the observed behavioral effects. Overall brain dynamics highlight reciprocal influences of working memory and selective attention processes, in keeping with shared cognitive resources between them.

## Introduction

Attention and working memory are constantly at play in cognitive tasks and have been the subject of intense experimental work. Yet only a fraction of these studies examines the interplay between these two major cognitive processes. We focus here on auditory selective attention, working memory, and their interactions as revealed by brain imaging studies.

### Auditory selective attention and working memory: brain networks

Numerous studies have shown that auditory selective attention and working memory rely on distributed brain networks, encompassing auditory temporal areas, prefrontal, and parietal cortex. On the one hand, using fMRI or PET, auditory attention has been shown to enhance activation in the auditory cortices in the superior temporal gyri (e.g., O’Leary et al., 1997; Lipschutz et al., 2002; Petkov et al., 2004). Similarly to the visual modality, top-down auditory attention is found to be supported by the dorsal frontoparietal network of attention (dorsal attention network, DAN), including the frontal eye fields, the inferior and middle frontal gyri, the anterior cingulate cortices, and parietal regions, with greater activation in the right hemisphere (e.g., Lipschutz et al., 2002; Kawashima et al., 1999; Belin et al., 1998). However, in the case of auditory spatial selective attention (attention to left or right ear), activation of this DAN and of the auditory cortices is greater contralaterally to the to-be-attended side (e.g., Lipschutz et al., 2002; Tzourio et al., 1997; Alho et al., 1999). On the other hand, auditory working memory networks have also been studied with fMRI and PET, mostly using verbal stimuli (for a review see Buchsbaum & D’Esposito, 2019) but also with melodic (tonal) material (e.g., Zatorre et al., 1994; Schulze et al., 2011). During sound sequence encoding, these neuroimaging studies have shown that activity increases in auditory cortices and the prefrontal cortex, with a left-hemisphere bias especially in prefrontal areas for verbal stimuli (meta-analysis in Buchsbaum et al., 2011), and more bilateral activity patterns for tonal stimuli (Gaab et al., 2003; Albouy et al., 2019). The maintenance (rehearsal) of auditory information recruits prefrontal areas, especially in the left hemisphere for verbal stimuli (e.g., Buchsbaum et al., 2011; Albouy et al., 2019) and bilaterally for tonal stimuli (Albouy et al., 2019), but also other areas including the hippocampus (Kumar et al., 2016), premotor and motor areas (Koelsch et al., 2009; Schulze et al., 2011), cerebellum (Gaab et al., 2003), and the Sylvian-parietal-temporal (Spt) area for verbal rehearsal (meta-analysis in Buchsbaum et al., 2011). The Intra-Parietal Sulcus (IPS) is involved in tonal working memory, especially when manipulation of the stored information is requested (Foster & Zatorre, 2010a, 2010b).

Neuroimaging studies that directly compare the networks subtending selective attention and working memory are scarce, especially in the auditory modality. A notable exception is the study by Huang et al. (2013), where they manipulated the level of attentional interference and memory load in an auditory Continuous Performance Task. Both manipulations resulted in activity changes in a fronto-parietal network, which were largely overlapping, with however some regions being sensitive only to memory load (notably in the anterior dorsolateral prefrontal cortex) and others only to attentional interference (notably in the Inferior Frontal Cortex).

### Auditory selective attention and working memory: network dynamics

The brain dynamics of auditory selective attention has been extensively investigated using non-invasive EEG and MEG and the dichotic paradigm introduced by Hillyard and collaborators (Hillyard et al., 1973). By comparing the evoked responses to the same sounds when task-relevant or irrelevant, attention has been shown to enhance transient responses as early as 20ms after sound onset (e.g., Hillyard et al., 1973; Woldorff et al., 1987; Woods et al., 1991; see Giard et al., 2000 for a review) and sustained responses (Picton et al., 1978; Bidet-Caulet et al., 2007). Importantly, recordings directly from the auditory cortices using intracortical EEG in epileptic patients have shown that auditory selective attention is supported by both an enhancement of evoked responses to relevant sounds and a suppression of evoked responses to irrelevant information, in primary and secondary auditory areas (Bidet-Caulet et al., 2007). It suggests that both facilitatory and inhibitory mechanisms are at play (see also Michie et al., 1993; Alho et al., 1987, 1994; Bidet-Caulet et al., 2010; Chait et al., 2010; Donald, 1987; Schroger & Eimer, 1997). They are modulated by task difficulty: on one hand, the difficulty of the task at hand (e.g., more difficult discrimination of to-be-attended sounds) seems to increase the brain responses to to-be-attended sounds while not affecting the brain responses to to-be-ignored sounds (Lange & Schnuerch, 2014). On the other hand, the difficulty of distractor filtering was found to reduce the processing of to-be-attended sounds (Alain & Woods, 1994; Alain et al., 1993; Ponjavic-Conte et al., 2013) or to increase the brain responses to to-be-ignored sounds (Melara et al., 2005). Alpha oscillations have also been found to be involved in top-down auditory attention. During anticipation and processing of a target sound, the power of alpha oscillations decreases in task-relevant auditory regions and increases in task-irrelevant occipital regions (Müller & Weisz, 2012; Weisz et al., 2014; Frey et al., 2014; Weise et al., 2016; ElShafei et al., 2018, 2020; Wöstmann et al., 2016), in line with an inhibitory role of the alpha rhythm (e.g., Klimesch et al., 2007; Foxe & Snyder, 2011; Jensen & Mazaheri, 2010; Thut et al., 2006; Palva & Palva, 2011). Interestingly, distinct frequency peaks in the alpha band have been found for the power increase in the occipital regions (around 11 Hz) and the power decrease in the auditory regions (around 9 Hz) (ElShafei et al., 2018, 2020), emphasizing the importance of considering the alpha frequency peak in different brain regions (see also Haegens et al., 2014).

EEG and MEG studies of auditory working memory have revealed a modulation of evoked responses by memory load and specific oscillatory signatures of working memory processes in various frequency bands. The amplitude of the N1 evoked response is related to memory performance: the N1 is reduced (and delayed) during encoding of short melodies in participants with congenital amusia who have a specific deficit of pitch short-term memory (Albouy et al., 2013). When memory load increases, the amplitude of sustained evoked responses during the maintenance in memory increases, with a specific component recorded over frontal areas reflecting auditory maintenance, the Sustained Anterior Negativity (SAN, e.g., Lefebvre et al., 2013; Grimault et al., 2014). Sustained evoked responses are also observed during the encoding of sound sequences, showing an increase in amplitude when the difficulty of the memory task increases (Albouy et al., 2013), which is very similar to the effect of attention observed during sound sequence processing (Bidet-Caulet et al., 2007). Oscillatory activities have been largely studied during working memory tasks, in particular during the maintenance period, when there is no actual stimulus presented. In a recent systematic review, Pavlov and Kotchoubey (2022) highlighted that the most consistent findings were an increase of frontal midline theta oscillations during both verbal and visuo-spatial working memory tasks, as well as an occipito-parietal alpha increase during verbal memory tasks. A majority of studies report that frontal theta oscillations increase with memory load, and these oscillations have been linked with the maintenance of items’ order (e.g., Hsieh et al., 2011), through a coupling with higher-frequency (gamma) activity (Canolty et al., 2006; Lisman & Jensen, 2013). Regarding alpha oscillations, the occipito-parietal alpha power increases with increasing memory load during verbal short-term memory tasks (Obleser et al., 2012) and could reflect the inhibition of the task-irrelevant visual areas (Klimesch et al., 2007), to protect the memory traces. EEG and MEG studies of oscillatory activities during working memory for non-verbal auditory material are far less numerous than studies about verbal working memory. However, for example during a melody short-term memory task, an increase of alpha power in task-irrelevant regions, namely the left fronto-temporal pathway and bilateral visual areas, has been observed (see Tillmann et al., 2016), suggesting similar principles of organization across auditory verbal and non-verbal domains.

### Interactions between selective attention and working memory

The studies reviewed above suggest: (1) largely overlapping brain networks involved in auditory selective attention and working memory, and (2) some similarities in how filtering out distractors and memory load impact on brain network dynamics. In particular, sustained evoked responses increase when selecting the relevant information in attention tasks and when memory load increases in working memory tasks. In both attention and working memory tasks, alpha oscillations increase in non-relevant brain areas. Some studies have tested more directly interactions between selective attention and working memory.

Such interactions have been studied extensively in the visual modality. One prominent finding is increased distraction effects at the behavioral and cerebral levels (increased responses to distractors) under high memory load in dual task-paradigms (one WM task and one attention task), in agreement with the Cognitive Load theory (de Fockert et al., 2001; Lavie et al., 2004; Lavie, 2005). However, in studies using single tasks (one WM task with attentional filtering of distractors), reduced distraction has been found under high memory load (Sörqvist, Stenfelt, et al., 2012; Sörqvist, Nöstl, et al., 2012; Sörqvist & Marsh, 2015). Similarly, in the few studies exploring the effect of memory load on auditory selective attention, inconsistent results have been found, with evidence for both enhanced (Bidet-Caulet et al., 2010) or reduced distraction effects (Bayramova et al., 2021; Berti & Schröger, 2003; SanMiguel et al., 2008) with increasing memory load. The other way around, auditory distraction stemming from irrelevant speech or non-speech sounds has been shown to impede on WM performance, mostly assessed with visually-presented verbal material (e.g., Salamé & Baddeley, 1989; review in Banbury et al., 2001). Interestingly, modulations of both memory load and attentional filtering were found to be reflected in activity patterns in the DAN using fMRI (Majerus et al., 2018).

Such studies either manipulated memory load and assessed its impact on distractors processing, or manipulated filtering difficulty and assessed its impact on memory processes. A joint investigation of both types of manipulation, together with neurophysiological measurements, would however be needed to further understand the interplay between working memory and selective attention.

### The current study

In the current study, to shed light on the links between selective attention and working memory networks, we take advantage of a newly designed paradigm, MEMAT (Blain, Talamini, et al., 2022), which allows to manipulate both selective attention and working memory processes during sound sequence encoding and rehearsal, using melodies (hence non-verbal auditory material). In MEMAT, participants perform a delayed-matching-to-sample task (DMST) using melodies presented in one (cued) ear. They have to indicate whether two consecutive sequences of four notes, separated by a two-second silent retention delay, are identical or not. During encoding of the first sequence, an interleaved to-be-ignored melody is presented in the other ear. The to-be-ignored melody can be more or less similar to the to-be-attended melody, hence allowing to manipulate the difficulty to filter out distractors (selective attention manipulation), and the memory task require more or less precision during encoding, depending on blocks, thus allowing to manipulate the memory load (see Blain, Talamini, et al., 2022). In previous reports (Blain, de la Chapelle, et al., 2022; Blain, Talamini, et al., 2022), we have shown that behavioral results obtained with the MEMAT paradigm are in favor of some shared cognitive resources between working memory and selective attention in the auditory non-verbal domain, which can be interpreted within the Cognitive Load theory framework (Lavie et al., 2004; Lavie, 2005), i.e., inhibition of distractors would be easier under low memory load. However the results also point to differences between working memory and selective attention processes, as revealed by slightly different patterns of results in various participants groups: on the one hand, musicians exhibit greatly enhanced working memory abilities for musical sequences but only limited advantages if any for selective attention compared to non-musicians (Blain, Talamini, et al., 2022), and on the other hand, participants with high dream recall frequency are more sensitive to the presence of difficult-to-filter-out distractors than participants with a low dream recall frequency, whereas the impact of the memory task difficulty manipulation was similar is these two participant groups (Blain, de la Chapelle, et al., 2022). These between-groups differences reveal that attention and working memory are differently sensitive to expertise effects (for the effect of musical expertise on auditory working memory see for example George & Coch, 2011; Talamini et al., 2016, 2017, 2022) and psychophysiological traits (for the effect of such traits on auditory attention, see Ruby et al., 2013; Eichenlaub, Bertrand, et al., 2014; Eichenlaub, Nicolas, et al., 2014; Vallat et al., 2020, 2022; Ruby et al., 2022). Overall, behavioral results obtained so far with the MEMAT paradigm revealed both shared and separated processes subtending auditory selective attention and working memory. Here we use MEG recordings during performance of MEMAT to tackle the interplay between selective attention and working memory at the neurophysiological level, using non-verbal auditory sequences. Based on the literature reviewed above, we focus our investigation on both transient and sustained evoked responses, as well as on oscillatory activities in the theta and alpha range, in auditory, visual, and frontal areas.

## Methods

The experimental protocol is adapted from the recent behavioral-only MEMAT study (Blain, Talamini, et al., 2022). The main difference is that the current data is based on a much larger number of trials, to achieve a good signal-to-noise ratio in the MEG data analysis.

### Participants

Twenty-two non-musician participants (20 right-handed, 11 men and 11 women, aged 20-33 years, mean education level ± standard deviation, SD: 15.5 ± 2.0 years) participated in the experiment. Participants were considered as non-musicians when they practiced less than 1 year of instrument or singing outside compulsory educative programs (mean music education level ± SD: 0.4 ± 0.7 years). Note that these participants were selected to have low or high frequency of dream recall, in order to be able to study the relationship between dream recall frequency (DRF), attention, and short-term memory in analyses not presented here. Behavioral data collected in the present study were analyzed as a function of participants’ DRF in Blain, de la Chapelle et al. (2022). Detailed neuropsychological testing and questionnaire data can be found in Blain, de la Chapelle et al. (2022). The present report focuses on MEG data combined from all participants.

All participants were free from neurological or psychiatric disorder and had normal hearing and normal vision. All subjects gave written informed consent to participate. Experimental procedures were approved by the appropriate regional ethics committee.

### Stimuli

240 four-sound-long melodies were created to be used as S1 melodies (see below) thanks to combinations of eight-harmonic synthetic sounds from the C major scale, spanning four octaves between 65 and 1046Hz. The maximal interval between any two sounds of a melody is 7 semi-tones. Sounds are 250-ms long and the interval between sounds (offset to onset) is 250 milliseconds. In each melody, there are no consecutive identical sounds. Every melody contains at least an ascending and a descending interval.

### Paradigm

#### Attention manipulation

The to-be-attended melody (S1) and the to-be-ignored melody (DIS) are played one in each ear. The tones of the two melodies are interleaved, thus not played simultaneously. S1 is played in the ear indicated by an arrow on the screen, and DIS is played in the other ear, each tone one after the other. The first sound to be played is the first tone of S1, and the last tone to be played is the last tone of DIS (see Figure 1).

**Figure 1.**
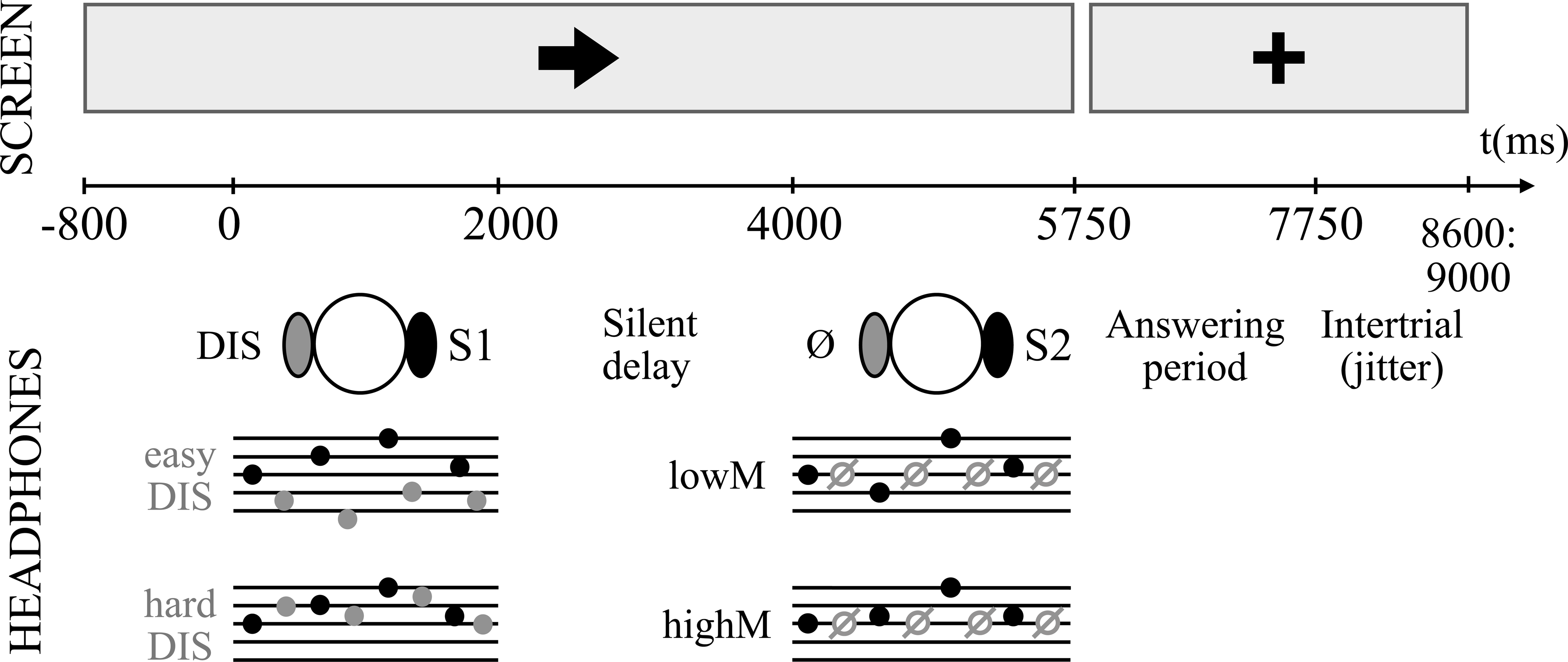
Trial design. The same paradigm is described in Blain, de la Chapelle et al. (2022). The ear of interest is indicated by an arrow on the screen during the trial. The to-be-attended melody S1 played in this ear must be encoded in memory while a to-be-ignored melody DIS is played in the other ear, interleaved with S1 tones. After a 2s retention delay, a melody S2 is played in the ear of interest and participants indicate whether S1 and S2 are identical or different. The melody DIS can be either easy (easyDIS) or hard (hardDIS) to ignore based on their frequency similarity with S1. When S2 is different from S1, one note is changed of either 5-6 semi-tones, entailing a contour change in the melody (low difficulty memory task; lowM blocks) or 1-2 semi-tones without contour change (high difficulty memory task; highM blocks).

The DIS melodies can be rather easy (easyDIS) or hard (hardDIS) to filter. An easyDIS melody is composed of sounds in a frequency range 6 to 7 semi-tones higher (or lower) than the corresponding S1 melody frequency range. A hardDIS melody frequency range is the same as the corresponding S1 melody frequency range. DIS melodies are constructed with the same rules as for the S1 melodies, as described above. Presentation of easy- and hard-to-ignore distractors is randomized within a given block therefore participants are not aware in advance of their difficulty.

#### Memory task

For each trial (see Figure 1), the participant is informed of the ear of interest by an arrow (all visual contents are seven-centimeter-wide black elements displayed individually at the center of a white screen). 800ms after the arrow presentation onset, S1 and DIS are played in the ear of interest and in the other ear, respectively. After a 2000-ms pause, the melody (S2) to compare to S1 is played in the ear of interest. At the end of S2 presentation, the indicative arrow is replaced by a central fixation cross. The participant has then 2000 ms to answer the question “Is S2 identical or different from S1?” using two response buttons (index and middle fingers of the right hand for identical and different respectively). When S2 differs from S1, only one sound out of four differs, and it can be either the second, third, or fourth one, never the first one which would be too salient. The answer period starts at the end of the S2 melody. At the end of the 2-second-long answering period, there is a pause lasting randomly from 850 ms to 1250 ms before the start of the next trial.

In the low memory task (lowM), when S2 differs from S1, one sound is replaced by another which is 5 to 6 semi-tones apart (within the C-major scale, ascending or descending) and induces a change in the melody contour. In the high memory task (highM), when S2 differs from S1, the changed sound differs from the original one only by 1 or 2 semi-tones (within the C-major scale) and does not induce a change in the melody contour. The difficulty of the memory task (MEMdiff) is fixed within a block, participants are notified of this difficulty at the beginning of each block.

#### Procedure

First, participants fill in neuropsychological and demographical tests and questionnaires. Then, they are installed in the MEG in a seated position, in a sound-attenuated, magnetically-shielded recording room, at 50-cm distance from the screen. All stimuli are delivered using Presentation software (Neurobehavioural Systems, Albany, CA, USA). Sounds are delivered through air-conducting plastic ear tubes. Detection thresholds for each ear are measured with an adaptive tracking procedure for A3 (220Hz, which is at the middle of the range of notes used). The auditory stimuli are subsequently presented at a comfortable listening level (60 dB SL). Then brain activity is recorded during a 5-min resting state period (eyes opened with a fixation cross provided to the participant). Participants then perform the main task. They are instructed to answer as accurately as possible with their right hand and to keep fixating the cross during the recordings. Prior to the actual task, they perform two short training blocks (12 trials) for each memory difficulty level. Training melodies (S1, DIS, and S2) are specific to the training sessions. Finally, they perform the ten experimental blocks (48 trials each), 5 under each memory difficulty (low or high). They are aware in advance of the difficulty of the memory task but not of the difficulty of the distractors, which is randomized within each block. The order of the blocks is randomized between participants. Trials last 9400 to 9800ms, leading to eight-minute-long blocks.

#### Balancing – within blocks

Each of the ten blocks contains 48 trials. All the combinations of ear of interest (Left or Right), distractor difficulty (easyDIS or hardDIS), associated answer (same or different) are equiprobable within block (6 trials/block each). For each combination with an expected answer “different”, the direction of the change is equiprobably ascending or descending in frequency (3 trials/block each). This within-block balancing allows to control for side, expected answer, and change direction.

#### Balancing – between blocks

The same S1 sequences are used in lowM and highM. For half of the participants in each group, melodies S1 and DIS are inverted and become respectively DIS and S1. Association between one S1 melody and another DIS melody changes across participants, enabling us to limit the impact in the results from any unfortunate easy or difficult match.

### Behavioral Data

#### Trial rejection

Trials in which participants gave no answer, several answers, or did answer at an inappropriate moment (*i.e.,* outside of the indicated period) are excluded from analysis. Average number ± SD of trials excluded per participant is 4 trials ± 4 (out of 480).

#### Measurements

Data are analyzed using Signal Detection Theory measures. dprime (d’) is the difference between normalized “Hits” and normalized “False alarms”. Hits are the proportion of correct “different” answer over all “different” trials, and False alarms are the proportion of incorrect “different” answer over all “same” trials. Criterion (c) indicates the response bias. Positive values indicate the tendency of answering “same” and negative values reflect the tendency of answering “different”. Median Reaction Times (RTs) are computed out of correctly answered trials and correspond to the time between the end of the S2 and the button press.

#### Statistical analyses for the behavioral data

All analyses are conducted using Bayesian statistics, which allow to test the similarity between measures and to estimate a degree of logical support or belief regarding specific results, as implemented in the JASP software (v0.16.3, JASP Team, 2022; Marsman & Wagenmakers, 2017; Wagenmakers et al., 2018; van den Bergh et al., 2022). We report Bayes Factor (BF_10_) as a relative measure of evidence (compared to the null model). To interpret the strength of evidence against the null model, we consider a BF between 1 and 3 as weak evidence, a BF between 3 and 10 as positive evidence, a BF between 10 and 100 as strong evidence, and a BF higher than 100 as a decisive evidence (Lee & Wagenmakers, 2014). Similarly, to interpret the strength of evidence in favor of the null model, we consider a BF between 0.33 and 1 as weak evidence, a BF between 0.01 and 0.33 as positive evidence, a BF between 0.001 and 0.01 as strong evidence and a BF lower than 0.001 as decisive evidence. For clarity, we report information about the best model only, unless several models account almost as well for the data (as assessed when comparing models to the best model). Additionally, we report the results of an analysis of effects that compares models that contain a given effect (of a factor or an interaction) to equivalent models stripped of the effect and was considered as a relative measure of evidence supporting the inclusion of a factor (BFinclusion).

d’, criterion (c), and RTs are submitted to a Bayesian repeated-measure ANOVA with MEMdiff (difficulty of the task, two levels: lowM, highM), DISdiff (difficulty of the DIStractor, two levels: easyDIS, hardDIS) as within-participant factors. Post-hoc comparisons for main effects or interactions are conducted using Bayesian t-tests. Criterions (c) for the four conditions (2 levels of MEMdiff x 2 levels of DISdiff) are submitted to a Bayesian one-sample t-test comparison to zero.

### Brain recordings

MEG data are continuously recorded using a 275-channel whole-head axial gradiometer system (CTF-275 by VSM MedTech Inc., Vancouver, Canada) with a sampling rate of 600Hz and a 0.016-150Hz bandpass filter. Head movements are continuously monitored using 3 coils placed at the nasion and the two preauricular points.

For each participant, a T1-weighted 3D MRI is obtained using a 1.5T or 3T whole-body scanner (Magnetom Siemens Erlangen, Germany) after the MEG session. At the end of MEG session, locations of the nasion and the two preauricular points are marked using fiducials markers visible on the T1 acquisitions. The T1 images are used for reconstruction of individual head shapes to create forward models for the source reconstruction procedures.

### Data pre-processing

MEG data are pre-processed offline using the software package for electrophysiological analysis (ELAN Pack) developed at the Lyon Neuroscience Research Center (Aguera et al., 2011). A third-order spatial gradient noise cancellation is applied to the data. The recorded signal is rejected when head movements exceed the threshold of +/-1cm around the median position. Sensor jumps (> 10 picoTesla) are detected and corrected with custom software. Blinks, saccades, and heartbeats are removed from the signal thanks to an Independent Component Analysis (ICA), using eeglab MATLAB toolbox (Delorme & Makeig, 2004). This ICA is computed on band-pass filtered signal (0.5-45Hz). Eye-movements and heartbeat-related components are determined by visual inspection of component topographies and time courses and removed through an ICA inverse transformation applied to the non-filtered signal. The ICA-corrected signals are then band-stop-filtered between 47 and 53 Hz, 97 and 103 Hz, and 147 and 153 Hz (zero-phase shift Butterworth filter, order 3) to remove power-line artifacts. Data segments contaminated with any residual artifact are excluded using a threshold of 2.5 picoTesla. 92% (±11) of trials remain in the analyses after artifact rejection (resulting in 55 (±6) trials per presentation side of S1 and condition).

### ERF analyses

#### ERF at sensor-level

For each participant and condition, event-related fields (ERFs) are obtained by averaging band-stopped ICA-corrected MEG data locked to the onset of S1 (time-window: -1500 to 7000 ms, averaging epochs thus start 250 ms prior to cue onset, see Figure 2). To study the Contingent Negative Variation (CNV) after the visual cue, in anticipation of the melody to encode, ERFs are filtered with a 30-Hz low-pass filter and baseline corrected to the mean amplitude of the 250-ms period before cue onset (-1050 to -800 ms before S1 onset). To study transient evoked responses to tones of S1/DIS melodies and sustained responses during S1/DIS melodies or the retention delay, ERFs are filtered with a 2-30-Hz band-pass filter and with a 2-Hz low-pass filter, respectively. ERFs are then baseline corrected to the mean amplitude of the -250 to 0 ms period before S1 onset (see Figure 2 and Table 1).

**Figure 2.**
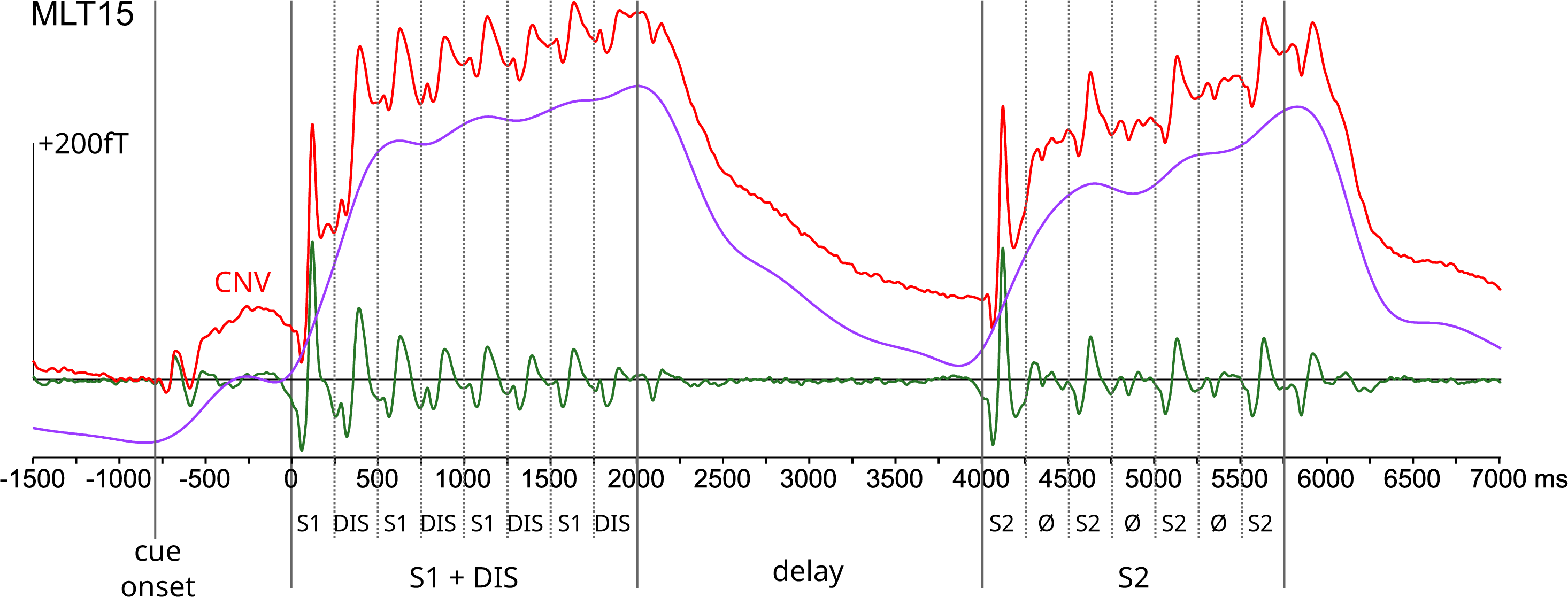
Filters used for ERF analyses (grand-average ERF). In red, a 30Hz low-pass filter was applied to preserve the CNV, between the onset of the visual cue (-800ms) and the onset of S1 (0ms), with a 1050 to -800 ms pre-cue baseline. In green, a 2-30Hz band-pass filter was applied to study the ERF to each S1 and DIS tone (between 0 and 2000ms), with a -250 to 0 ms pre-S1 baseline. In purple, a 2Hz low-pass filter was applied to highlight the sustained response during the S1/DIS melodies (0 to 2000ms) and the retention delay (2000 to 4000ms), with a -250 to 0 ms pre-S1 baseline.

**Table 1.**
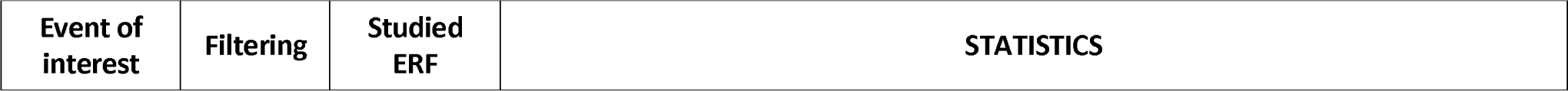
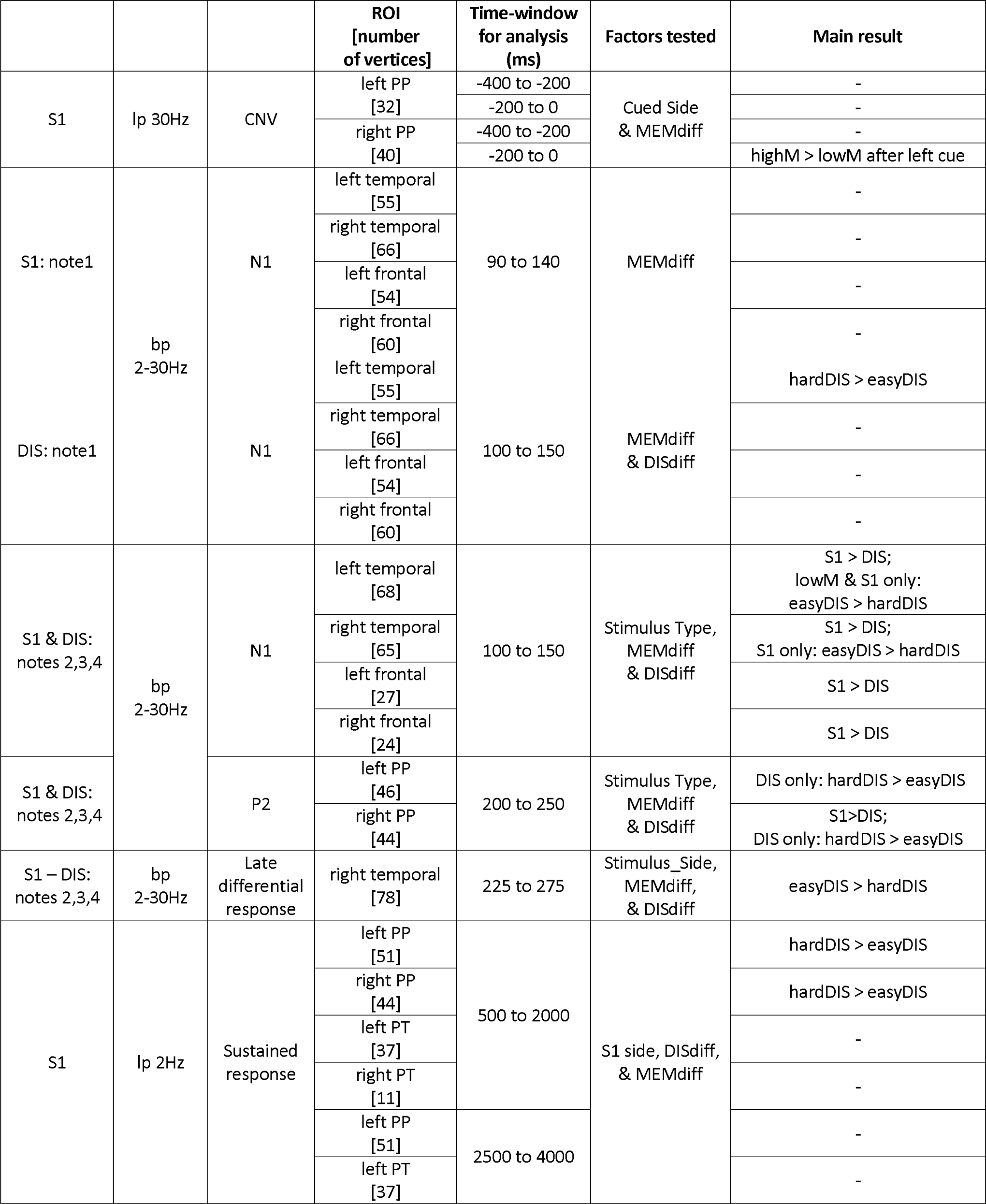
Summary of the ERF analyses: event and filtering used, studied ERF, ROI information, time-window used for statistical analysis, factors of the Bayesian repeated-measure ANOVA used, and results of the statistical analysis. For CNV and sustained responses, ROIs are defined based on activation from all stimuli between -400 and 0 ms and between 500 and 2000 ms respectively, whereas for N1, P2, and late responses, ROIs are defined based on activation from contralateral stimuli only (see text for details). When computing the difference between S1 and DIS, both S1 and DIS stimuli are contralateral (i.e. they are taken from different trials). For N1 and P2 for notes 2, 3, 4, as activity is largely overlapping for S1 and DIS, the ROIs are drawn from the emergence test of S1. We report in the table only effects for which at least positive evidence (i.e., BF_10_ or BF_inclusion_ > 3) was observed (see text for the full results). “>” means larger ERF (in absolute value) in a given condition compared to another one. “-” means no positive or greater evidence for an effect. S1: to-be-attended melody, DIS: to-be-ignored melody; lp: low pass; bp: band-pass; PP: Planum Polare; PT: Planum Temporale; MEMdiff: memory task difficulty (lowM vs. highM); DISdiff: distractor filtering difficulty (easyDIS vs. hardDIS).

Because of saliency effects for the first note, we further separate the analysis of the first note and the following ones: on the one hand, ERFs are computed locked to the first note onset, and on the other hand, we compute the average of the ERFs locked to the second, third, and fourth notes of the melodies, with no further baseline correction, separately for S1 and DIS in both cases. We also compute the difference between S1 and DIS (average across notes 2, 3, 4).

Grand-average ERF is obtained by averaging the same computed ERF across all 22 participants.

#### ERF source reconstruction

Segmentation of the individual T1-weighted MRIs is conducted using the FreeSurfer software package (Fischl, 2012; http://surfer.nmr.mgh.harvard.edu). Segmentation is visually inspected and then imported (15,002 vertices) in the Brainstorm toolbox (version 3.191209 (09-Dec-2019) on Matlab R2015a; Tadel et al., 2011; http://neuroimage.usc.edu/brainstorm) with which further source analyses are conducted. The white matter–grey matter boundary segmented by FreeSurfer is used as a source space for subsequent weighted minimum norm estimation. A noise covariance matrix is computed on resting state data, which has been preprocessed with the same pipeline as data collected during task performance. After coregistration between the individual anatomy and MEG sensors, the forward model is computed with the OpenMEEG software (Gramfort et al., 2010; https://openmeeg.github.io/). Cortical currents are estimated using a distributed model consisting of 15,002 currents dipoles from the time series of the 275 gradiometer signals, using a linear inverse estimator (weighted minimum-norm current estimate, signal-to-noise ratio of 3, whitening PCA, depth weighting of 0.5). Sources orientations are constrained to be orthogonal to the grey-white matter boundary of the individual MRIs. The results are then projected on a standard brain model anatomy (ICBM152; 15,002 vertices) for group analysis purposes.

#### ERF source analysis

ERF analyses are performed following five steps: (1) the main ERFs are identified at the sensor level, and time-windows of interest are defined for each ERF (see Table 1 for a list of ERFs studied and the associated time-windows); (2) at the source level, emergence of ERFs (all four conditions collapsed) is tested at the whole brain level using t-tests comparing signal amplitude (on average across the time-window of interest) to zero at each vertex (FDR-correction for multiple comparisons, corrected p<0.05); (3) regions of interest (ROIs) are drawn using Brainstorm based on these emergence tests with a procedure allowing to grow from a seed voxel to match the shape of the activation patch on the cortical surface; (4) ROI time-courses of activity are extracted for each participant, each condition, each presentation side; (5) mean amplitudes in time-windows of interest are computed for each participant, each condition, each presentation side; (6) Bayesian ANOVA (see Results and Table 1 for details about factors, and the Statistical methods for the behavioral data section above for details about Bayesian ANOVA) for each studied ROI and each time-window are performed (see Table 1 for ROIs and time-windows). Emergence of ERFs (step 2) is scrutinized for stimuli presented at left or right ear separately, and if similar patterns are observed, data are further collapsed across stimulus presentation side for ROI definition (see Table 1). Emergence of ERFs is also assessed in the difference between S1 and DIS (notes 2, 3, 4).

### Time-Frequency analysis

#### TF at sensor-level

To investigate the dynamics of theta and alpha power during the task, a time-frequency analysis is performed using the Fieldtrip toolbox (v20190626; Oostenveld et al., 2011) on Matlab R2017A. Time-frequency analysis is computed from -2500 to +8000ms relative to S1 onset, using Morlet wavelet decomposition with a width of four cycles per wavelet (m=7; Tallon-Baudry & Bertrand, 1999), for frequencies from 4 to 20Hz, with steps of 1Hz and 50ms. For each single trial, the corresponding ERF is subtracted before wavelet decomposition to focus on induced (and not evoked) activity. The relative change in power is then obtained by subtracting then dividing each power value with the power in a pre-cue baseline period (-1800 to -1000 ms relative to S1 onset) in the corresponding frequency range. The main power modulations are identified at the sensor level for all conditions collapsed and used to define time-frequency windows of interest for source-level analyses.

#### TF: head model

For source reconstruction, the processing of the T1 MRI is performed using CTF software (CTF Systems Inc., VSM Medtech Inc., Vancouver, Canada). For each participant, an anatomically realistic single shell headmodel based on the cortical surface is generated from individual head shapes (Nolte, 2003) and imported in the Fieldtrip toolbox (v20190626; Oostenveld et al., 2011). Then, a grid with 0.5-cm resolution is normalized on a MNI template and morphed into the brain volume of each participant.

#### TF: whole brain source analyses

To elucidate the possible brain regions underlying the sensor-level modulations in the theta and alpha bands, we perform whole brain analysis on time-frequency windows of interest.

Several time-windows relative to S1 onset are chosen based on the protocol design: one during the cue presentation (-800:0ms); two during the presentation of S1 and DIS melodies (400:1200ms & 1200:2000ms); and two during the delay (2400:3200ms & 3200:4000ms). Three different frequency bands (4 to 7Hz, 7 to 11Hz & 11 to 15 Hz) are chosen based on the observations at the sensor level and previous findings (ElShafei et al., 2018, 2020).

We use the frequency-domain adaptive spatial technique of Dynamical Imaging of Coherent Sources (DICS; Gross et al., 2001). For each participant, data from all conditions is concatenated, and the cross-spectral density (CSD) matrix (-2.5 to +7s, relative to S1 onset, lambda 5%) is calculated using the multitaper method with a target frequency of 9.5 (+/-5.5) Hz. Leadfields for all grid points along with the CSD matrix are used to compute a common spatial filter that is in turn used to estimate the spatial distribution of power for all time-frequency windows of interest.

Afterward, for left and right S1 presentation separately (irrespective of MEMdiff and DISdiff), power in each time-frequency window of interest is contrasted against the power in a corresponding baseline pre-cue window (-1800 to -1000 ms relative to S1 onset) for frequencies around 5.5 (+/- 1.5), 9 (+/- 2) and 13 (+/- 2) Hz using a nonparametric cluster-based permutation analysis (Maris & Oostenveld, 2007) across participants, controlling for multiple comparisons in the source space dimension.

#### Reconstruction of source-level single-trial activity in ROIs

In order to get a time-frequency resolved estimation of source activity, we use an approach based on regions of interest (ROIs) and we reconstruct single trial activity at the level of ROIs using the linearly constrained minimum variance (LCMV) beamformer (Van Veen et al., 1997) in each subject.

Regions of interest (ROIs), in each hemisphere, are defined from the Brodmann atlas based on the group whole brain activation (see previous paragraph): the *auditory ROI (Aud)* is composed of Brodmann areas 22, 41, and 42, the *visual ROI (Vis)* is composed of Brodmann areas 17, 18, and 19, the *anterior prefrontal cortex* ROI is composed of Brodmann areas 9 and 10, the *Broca ROI* is composed of Brodmann areas 44 and 45, and the *dorso-lateral prefrontal (DLPFC) ROI* is composed of Brodmann area 46.

Spatial filters are constructed from the covariance matrix of the averaged single trials at sensor level (-2.5:7s, relative to S1 onset, 1-20Hz, lambda 5%) and the respective leadfield for all grid points. Then, spatial filters are multiplied by the sensor-level single-trial data, thus obtaining the time course activity of each voxel of interest within the defined ROIs. Single-trial activity is averaged across all voxels within each ROI in each hemisphere. For each ROI, the evoked response (i.e., the signal averaged across all trials) is subtracted from each trial.

#### ROI peak frequency

To extract the frequency peak in each ROI (source-level activity reconstructed with LCMV, see previous paragraph), for each time-window of interest (-1800:-1000; -800:0; 400:1200; 1200:2000; 2400:3200; 3200:4000 ms), we compute the power spectral density using Fourier decomposition based on Hann windows and extract the rhythmic activity from the concurrent 1/f modulations of the signal using the Fitting oscillations and one-over-F (FOOOF) toolbox implemented in Fieldtrip (v20221004; Oostenveld et al., 2011; Donoghue et al., 2020), for all conditions collapsed. The resulting FFT of the baseline is subtracted from the resulting FFT of each following time-window.

The theta peak corresponds to the highest local maximum in the resulting spectrum between 3 and 7 Hz; while the peaks for alpha decrease and increase correspond to the lowest local minimum and highest local maximum between 7 and 15 Hz, respectively.

In each ROI, the group frequency range is then defined from the median peak frequency across subjects +/- 1.5 Hz, averaged over time-windows, resulting in 4-7 Hz, 8-11 Hz and 9- 12 Hz to be used in subsequent analyses for theta, low alpha and high alpha respectively.

#### ROI statistical analysis

In each ROI, the time-frequency power is calculated for each trial using Morlet wavelet decomposition with a width of four cycles per wavelet (m = 7) at frequencies between 4 and 15 Hz, in steps of 1 Hz and 50 ms. The relative change in power is then obtained by averaging single trial TF power in each condition (for each level of memory difficulty, distractor filtering difficulty, and side of S1 presentation) and by subtracting then dividing each averaged power value with the power in a pre-cue baseline period (-1800 to -1000 ms relative to S1 onset) in the corresponding frequency range.

For each frequency range (4-7 Hz, 9-12 Hz and 8-11 Hz) and each time-window of interest (-800:0; 400:1200; 1200-200; 2400:3200; 3200:4000 ms), the relative change in power is extracted in each subject and for each ROI of interest based on DICS analysis. To investigate the impact of the memory task and distractor difficulties, these averaged values are submitted to a Bayesian repeated-measure ANOVA with factors MEMdiff, DISdiff, and S1side, for each ROI and each corresponding time-frequency window (see Table 2). Post-hoc comparisons for main effects or interactions are conducted using Bayesian t-tests.

**Table 2.**
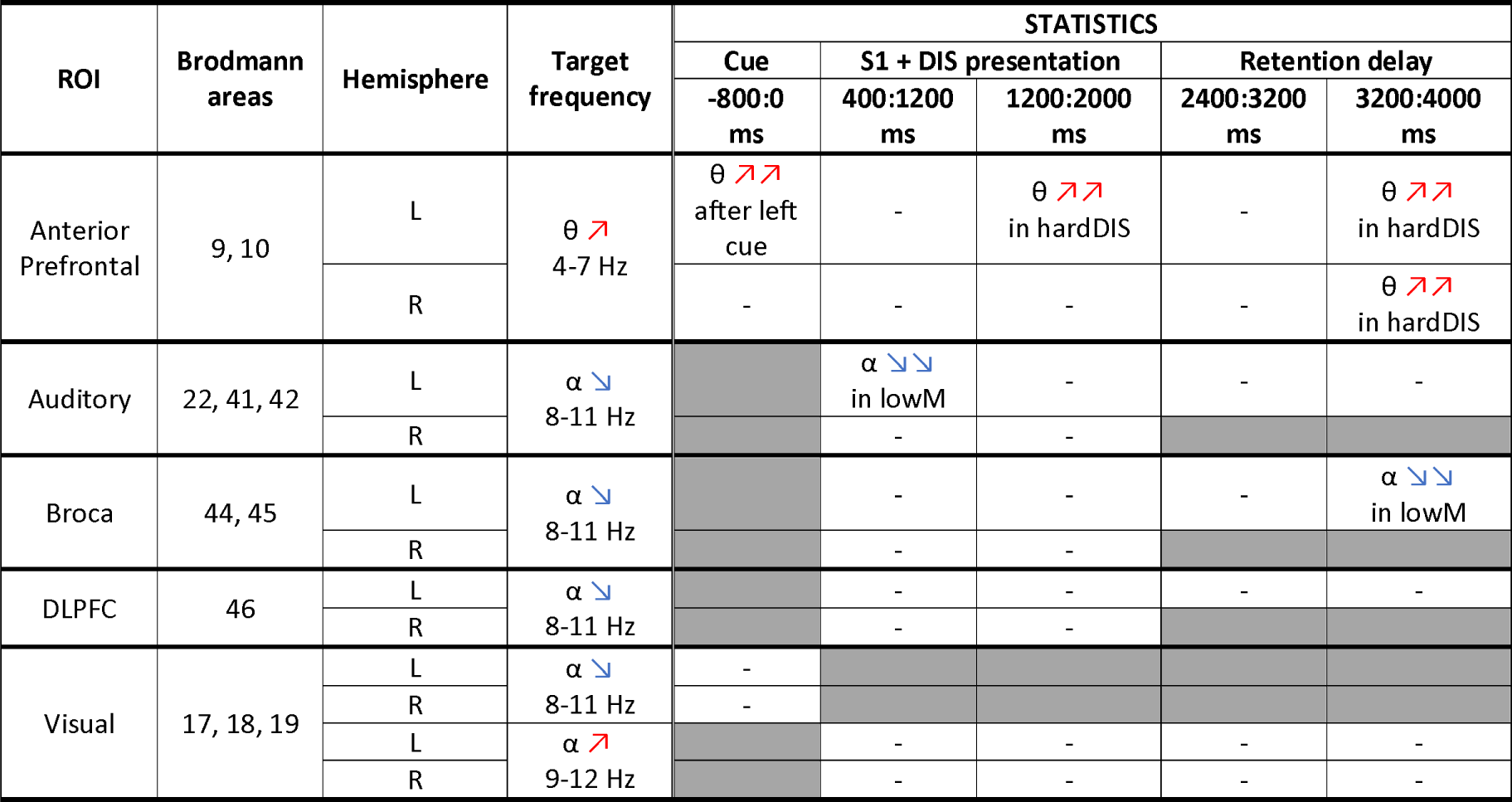
Summary of the TF analysis. We report in the table only effects for which at least positive evidence (i.e., BF_10_ or BF_inclusion_ > 3) was observed with a Bayesian repeated-measure ANOVA with factors S1side, DISdiff (distractor filtering difficulty, easyDIS vs. hardDIS), and MEMdiff (memory task difficulty, lowM vs. highM; see text for the full results). Reported effects are relative to the opposite condition. “-” means no positive or greater evidence for an effect. Gray cells were not tested in accordance with the whole-brain source results. ☐: increase in power, ☐: decrease in power.

| ROI | Brodmann areas | Hemisphere | Target frequency | STATISTICS |  |  |  |  |
| --- | --- | --- | --- | --- | --- | --- | --- | --- |
|  |  |  |  | Cue | S1 + DIS presentation |  | Retention delay |  |
|  |  |  |  | -800:0 ms | 400:1200 ms | 1200:2000 ms | 2400:3200 ms | 3200:4000 ms |
| Anterior Prefrontal | 9, 10 | L | $\theta$ ↗<br>4-7 Hz | $\theta$ ↗↗ after left cue | - | $\theta$ ↗↗ in hardDIS | - | $\theta$ ↗↗ in hardDIS |
| | | R | | - | - | - | - | $\theta$ ↗↗ in hardDIS |
| Auditory | 22, 41, 42 | L | $\alpha$ ↘<br>8-11 Hz | | $\alpha$ ↘↘ in lowM | - | - | - |
|  |  | R |  |  | - | - |  |  |
| Broca | 44, 45 | L | $\alpha$ ↘<br>8-11 Hz | | - | - | - | $\alpha$ ↘↘ in lowM |
|  |  | R |  |  | - | - |  |  |
| DLPFC | 46 | L | $\alpha$ ↘<br>8-11 Hz | | - | - | - | - |
|  |  | R |  |  | - | - |  |  |
| Visual | 17, 18, 19 | L | $\alpha$ ↘<br>8-11 Hz | - | | | | |
|  |  | R | 8-11 Hz | - |  |  |  |  |
| | | L | $\alpha$ ↗<br>9-12 Hz | | - | - | - | - |
|  |  | R | 9-12 Hz |  | - | - | - | - |

## Results

### Behavioral results

For d’ (Figure 3A), the best model explaining the data is the one with factors MEMdiff, DISdiff, and the interaction between MEMdiff and DISdiff (BF_10_=2.3e+16), in keeping with Blain, Talamini et al. (2022) and Blain, de la Chapelle et al. (2022). The analysis of effects across models reveals decisive evidence for MEMdiff (BF_inclusion_=3.7el+10), for DISdiff (BF_inclusion_=6729.8), and for the MEMdiff x DISdiff interaction (BF_inclusion_=116.0). d’ is higher for easyDIS compared to hardDIS, and higher in lowM compared to highM. The difference between hardDIS and easyDIS is smaller in highM compared to lowM (or put differently, the difference between lowM and highM is smaller for hardDIS than easyDIS), as revealed by a post-hoc Bayesian t-test (BF_10_=123.2).

**Figure 3.**
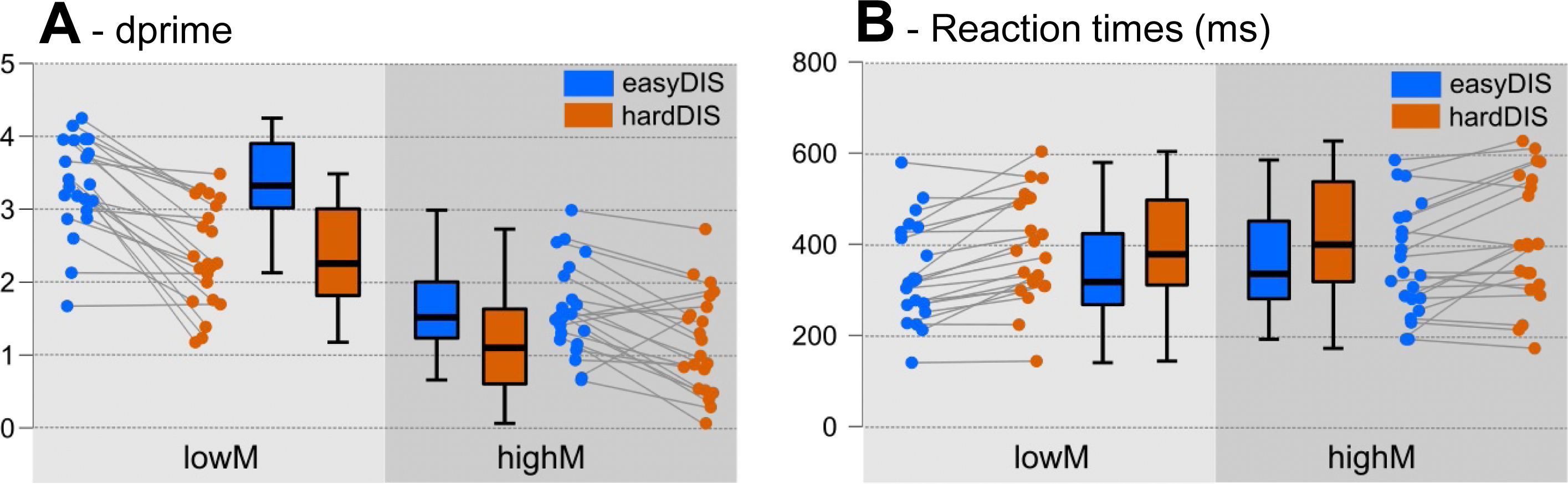
Behavioral results. Effect of the difficulty of the task (lowM: low difficulty, highM: high difficulty) and of the difficulty of the distractor (easyDIS: easy to ignore, hardDIS: hard to ignore) on performance d’ (A) and on reaction times (B). Whisker plots show the median±quartile and the minimum/maximum values. Lines in the strip plots connect data from the same participant.

For the criterion (c), in agreement with previous studies using delayed-matching-to-sample tests, there is weak to decisive evidence for positive c values in all conditions, underlining that participants tended to miss differences between melodies (see Table S5 in Blain, de la Chapelle, et al., 2022). The best model explaining criterion results is the one with factors MEMdiff, DISdiff, and the interaction between MEMdiff and DISdiff (BF_10_=4.4e+9). There is decisive evidence for MEMdiff (BF_inclusion_=1.4e+7), strong evidence for DISdiff (BF_inclusion_=73.7), and positive evidence for the MEMdiff x DISdiff interaction (BF_inclusion_=7.0). Criterion is higher under highM compared to lowM and is higher for easyDIS compared to hardDIS. Post-hoc Bayesian paired-samples t-tests reveals strong evidence for a difference in criterion between easyDIS and hardDIS for lowM (BF_10_=90.7) but only weak evidence for such a difference in highM (BF_10_=1.5).

For RTs (see Figure 3B), the best model explaining results is the one with factors MEMdiff and DISdiff (BF_10_=671.5). The analysis of effects reveals decisive evidence for DISdiff (BF_inclusion_=151.2) and positive evidence for MEMdiff (BF_inclusion_=4.6). RTs are longer for hardDIS compared to easyDIS, and they are longer for highM compared to lowM.

### Event-Related Fields Results

A summary of the main results obtained for ERF analyses is provided in Table 1.

#### Contingent Negative Variation (CNV)

The time course and topographies at the sensor level of the CNV developing after the visual cue are shown in Figure 4A. Emergence tests at the source level (time window: -400 to 0 ms relative to S1 onset) reveal activity in temporal regions before S1 onset, extending bilaterally along the Planum Polare, irrespective of the cued ear. Activity in these temporal regions was examined with Bayesian repeated-measure ANOVAs with Cued Side and MEMdiff as factors, during early (-400 to -200 ms relative to S1 onset) and late (-200 to 0 ms relative to S1 onset) CNV (see Table 1).

**Figure 4.**
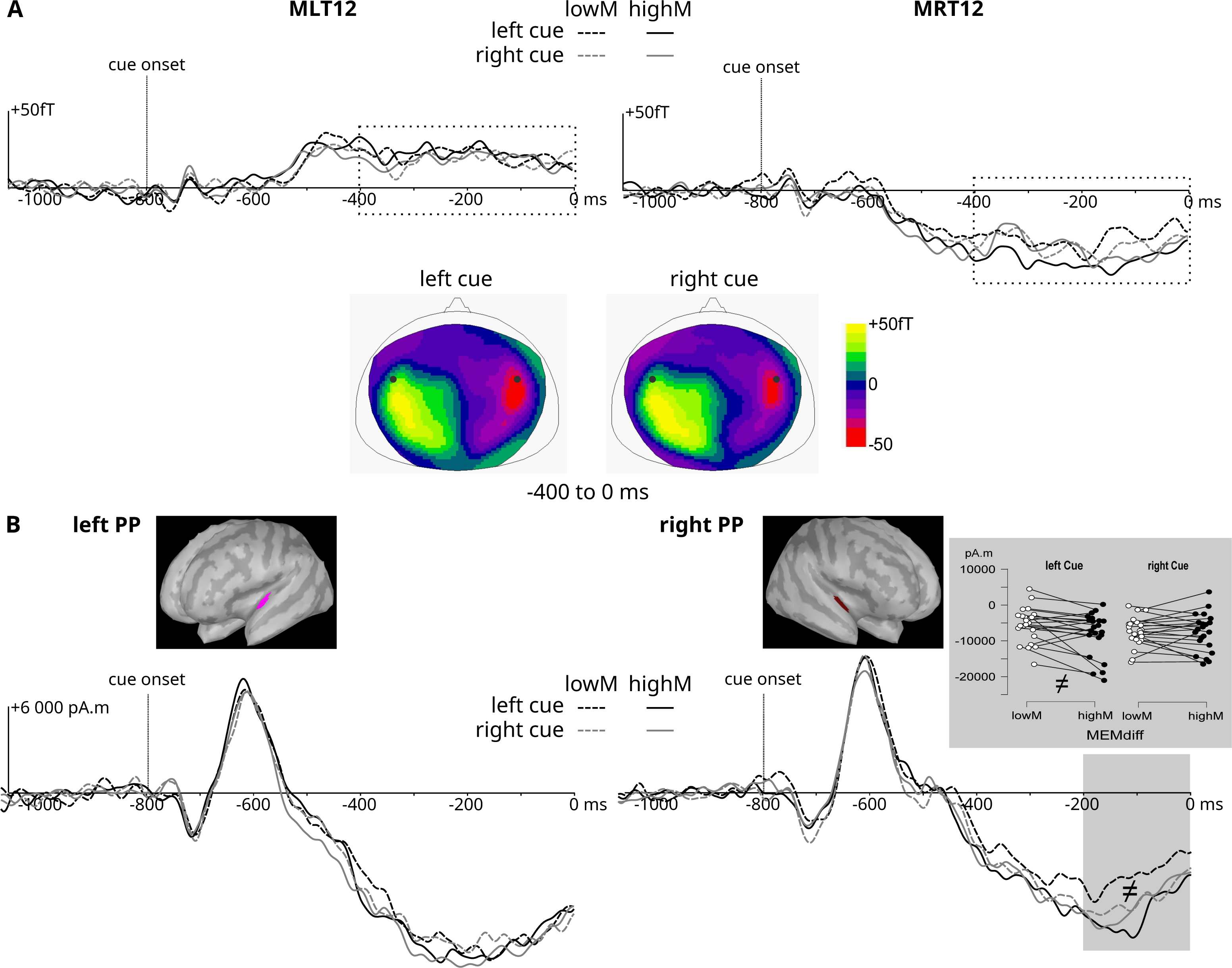
Contingent Negative Variation (CNV). (A) In the sensor space, time course of the ERF to the visual cue onset (arrow, -800ms relative to S1 onset), for each MEMdiff and Cued Side condition, at MLT12 and MRT12 (above the left and right temporal lobes, respectively). Topoplots show sensor data to left and right cues, over the time period of interest, indicated by the dotted rectangles (-400 to 0 ms). Black dots on topographies indicate corresponding sensor position. (B) In the source space, time course of the ERF to the visual cue onset, for each MEMdiff and Cued Side condition, in the corresponding ROI represented on the MNI template. The strip plot shows individual data averaged by condition in the gray period (-200 to 0 ms) showing positive or greater evidence (see Table 1 for time-windows and results). Lines connect data from the same participant. leftPP: left Planum Polare, right PP: right Planum Polare.

In the right Planum Polare ROI (see Figure 4B), the best model explaining the data just before S1 presentation (time-window -200 to 0 ms relative to stimulus onset) is the model with the MEMdiff factor (BF_10_ =1.2), immediately followed by the full model including the interaction between Cued Side and MEMdiff (BF_10_ =1.1). Note that these models are thus almost as likely as the null model. The analysis of effects reveals weak evidence for the interaction between MEMdiff and Cued Side (BF_inclusion_ = 2.6) and for MEMdiff (BF_inclusion_ = 1.2). Post-hoc Bayesian paired-sample t-tests revealed positive evidence for a difference between lowM and highM with contralateral stimuli (BF_10_ = 3.7) and for no difference between lowM and highM with ipsilateral stimuli (BF_10_ = 0.23). When the left ear was cued, larger CNV amplitudes are thus observed in the right Planum Polare for the highM condition compared to lowM.

The best model explaining CNV amplitudes in the left and right Planum Polare ROIs between -400 to -200 ms prior to S1 onset, as well as in the left Planum Polare ROI between - 200 and 0 ms prior to S1 onset, is the null model in all three cases (BF_10_ for all other models < 0.56).

25

#### Transient Evoked Responses: N1 to first tones of S1 and DIS

The time course of the transient evoked responses after each note in S1 and DIS melodies as recorded at the sensor level is shown in Figure 2. ERFs to the first notes of S1 and DIS are larger than ERFs to subsequent notes and are thus analyzed separately (see Figure 5). The first note of S1 and DIS evokes a large N1 response in contralateral temporal (Heschl Gyrus, HG, and around) and frontal (Inferior Frontal Gyrus, IFG) regions. Smaller responses with longer latencies are observed ipsilaterally. Therefore, the amplitude of the N1 is analyzed separately for each presentation side in the contra-lateral ROIs with Bayesian paired-value t-tests for the first note of S1 (S1-L and S1-R separately), to assess the effect of the MEMdiff factor, and with Bayesian repeated-measure ANOVAs with MEMdiff and DISdiff as factors for the first note of DIS (DIS-L and DIS-R separately).

**Figure 5.**
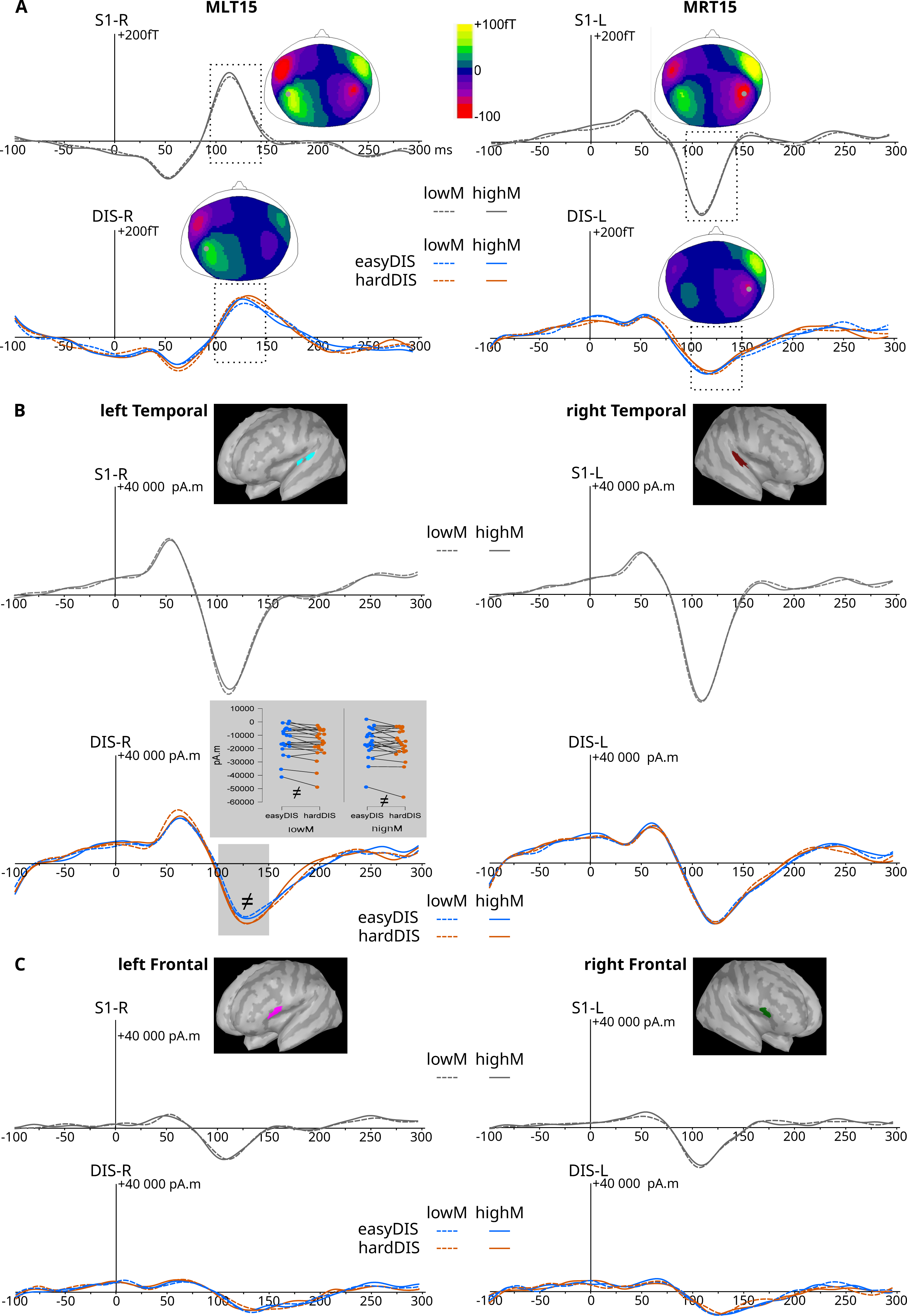
Transient evoked responses to the first tone. (A) In the sensor space, time course of the ERF to the first tone of the to-be-attended melody S1 and of the to-be-ignored melody DIS (tone onset at 0ms), for each DISdiff (DIS only) and MEMdiff condition (S1 and DIS), at MLT15 and MRT15 (above the left and right temporal lobes, respectively). Topoplots show sensor data over the time period of interest, indicated by the dotted rectangles (90 to 140 or 100 to 150 ms for S1 or DIS, respectively). Gray dots on topographies indicate corresponding sensor position. (B & C) In the source space, time course of the ERF to the first tone of S1 and DIS melodies, for each DISdiff (DIS only) and each MEMdiff condition, in the corresponding ROIs represented on the MNI template. The strip plot shows individual data averaged by condition in the gray period showing positive or greater evidence (see Table 1 for time-windows and results). Lines connect data from the same participant. Only contralateral tones are analyzed here: S1-R and DIS-R for left sensors and sources, and S1-L and DIS-L for right sensors and sources. S1-L or S1-R: S1 melodies presented to the left or right ear, respectively. DIS-L or DIS-R: DIS melodies presented to the left or right ear, respectively.

For the N1 elicited in response to the first presented tone (S1, time window 90-140 ms post S1 onset), no evidence is found for an effect of MEMdiff (all BF_10_ ≤ 0.65) in the four ROIs investigated (Table 1 and Figure 5B).

For the N1 elicited in response to the first to-be-ignored tone (DIS, time window 100-150 ms post DIS onset), the best model explaining the data in the left Temporal ROI (i.e., for DIS stimuli presented in the right ear) is the model with the DISdiff factor (BF_10_ = 3.6), followed by the model with DISdiff and MEMdiff (BF_10_ = 2.2), with the analysis of effects confirming only positive evidence for DISdiff (BF_inclusion_ = 3.7). A larger N1 is observed in response to the first DIS tone in hardDIS compared to easyDIS. In the left IFG ROI, the best model explaining the data (N1 to the first DIS tone) is the full model including the interaction between MEMdiff and DISdiff (BF_10_ = 1.9), however it is almost as likely as the null model and further Bayesian paired-value t-tests comparing N1 amplitudes across conditions reveal only weak evidence for differences between conditions, if any (all BF_10_ < 1.7). In the right hemisphere ROIs (temporal and IFG), for DIS stimuli presented in the left ear, the best models explaining the N1 amplitudes are the null model (for all other models, BF_10_ < 0.9).

#### Transient Evoked Responses: N1 to tones 2, 3, 4 (S1 and DIS)

Sensor-level ERFs for subsequent tones (S1 and DIS) are illustrated in Figure 6. Emergence tests reveal significant activity in contralateral temporal (HG and around) and frontal (IFG) regions at the latency of the N1 (100-150 ms post tone onset). Smaller responses with longer latencies are observed ipsilaterally. N1 amplitudes are analyzed in contralateral ROIs with Bayesian ANOVAs with StimulusType (S1 or DIS), MEMdiff, and DISdiff as factors (see Table 1 and Figure 7).

**Figure 6.**
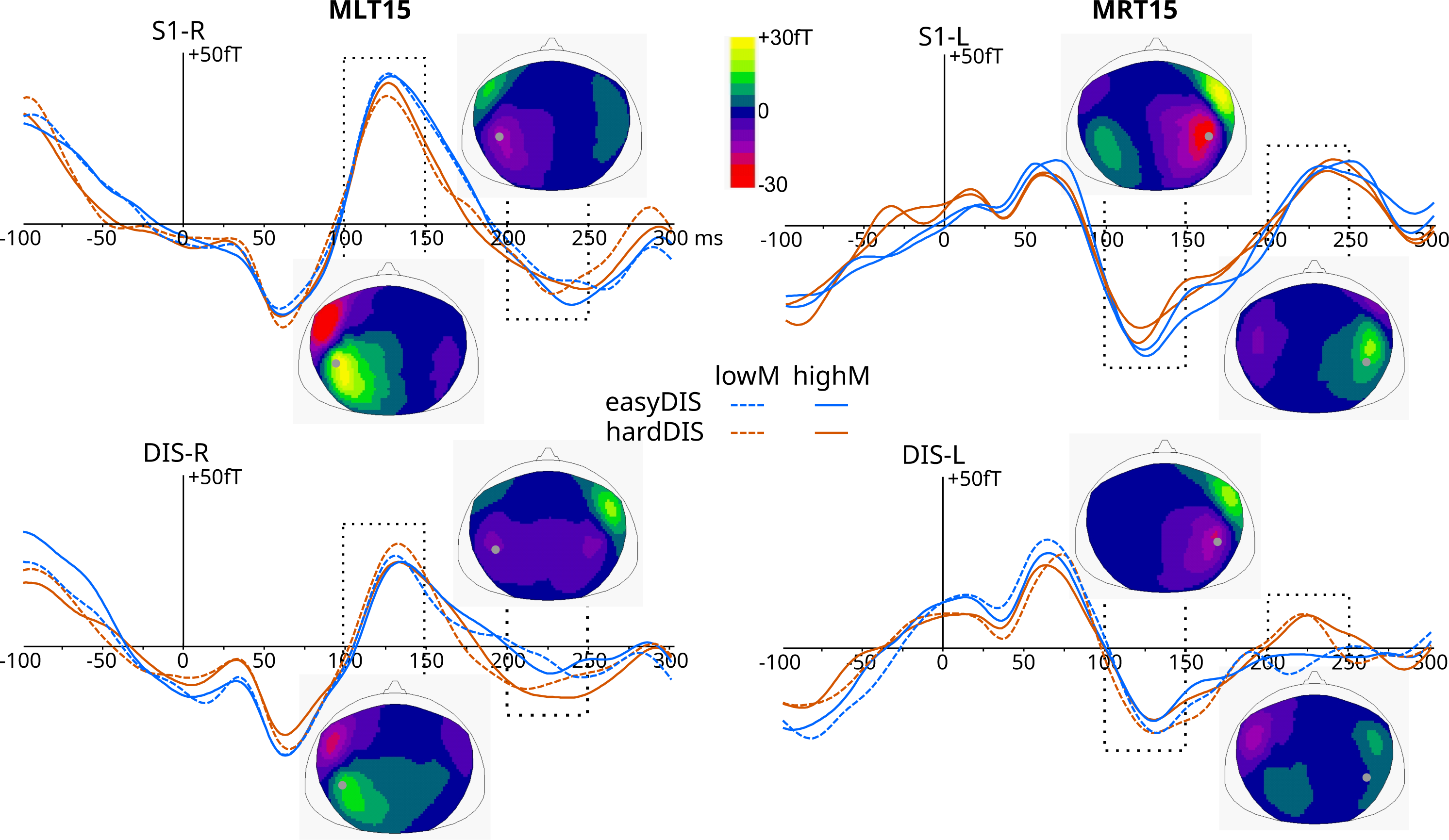
Transient evoked responses to the tones 2, 3 and 4 in the sensor space. Time course of the averaged ERF to the tones 2, 3 and 4 of the to-be-attended melody S1 and of the to-be-ignored melody DIS (tone onset at 0ms), for each DISdiff and MEMdiff condition, at MLT15 and MRT15 (above the left and right temporal lobes, respectively). Topoplots show sensor data over the time periods of interest, indicated by the dotted rectangles (100 to 150 ms and 200 to 250 ms for N1 or P2 responses, respectively). Gray dots on topographies indicate corresponding sensor position. Only contralateral tones are studied here: S1-R and DIS-R for left sensors, and S1-L and DIS-L for right sensors. S1-L or S1-R: S1 melodies presented to the left or right ear, respectively. DIS-L or DIS-R: DIS melodies presented to the left or right ear, respectively.

**Figure 7.**
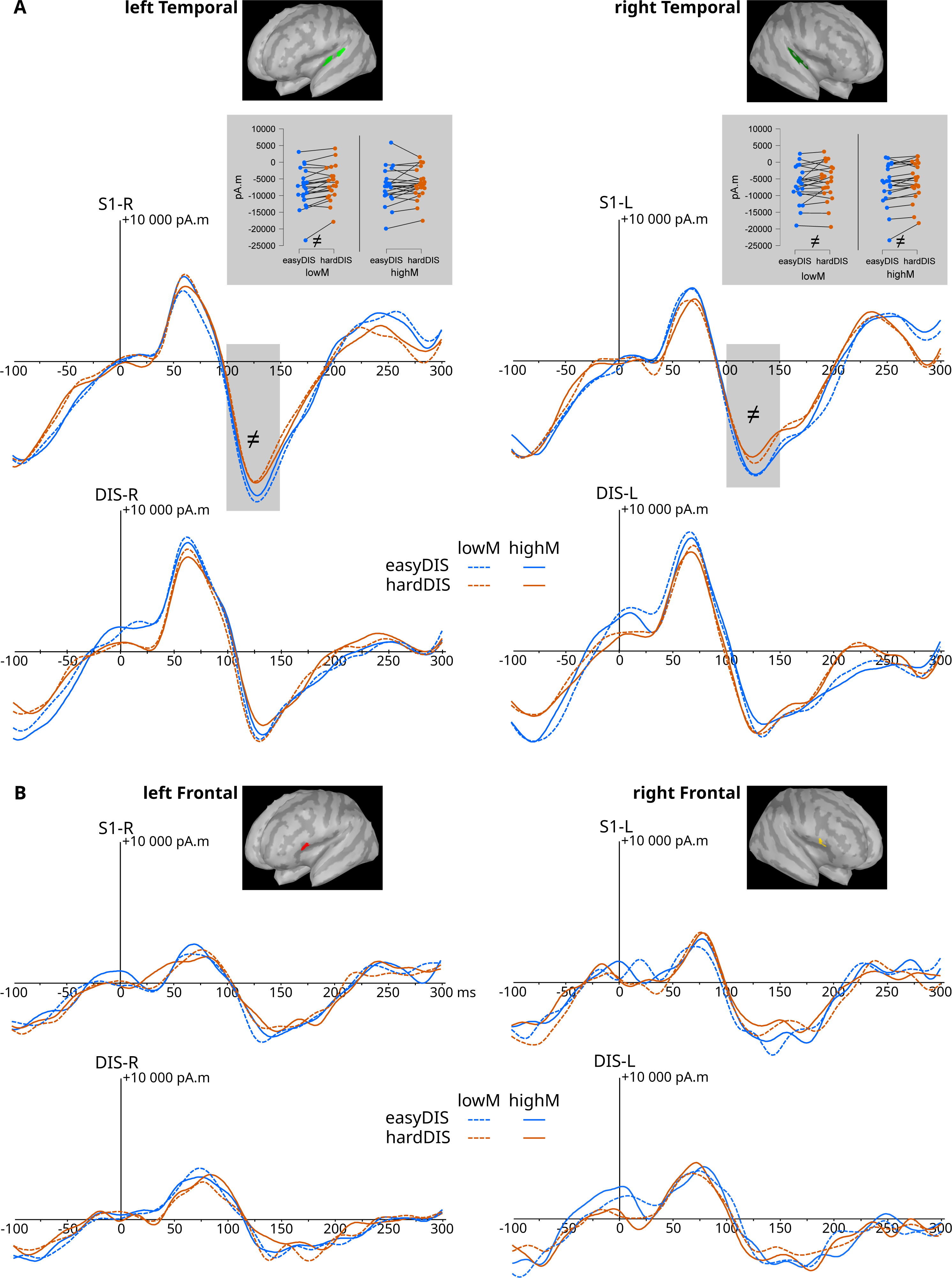
Transient evoked responses to the tones 2, 3 and 4 in the source space (N1 ROIs). Time course of the averaged ERF to the tones 2, 3 and 4 of the to-be-attended melody S1 and of the to-be-ignored melody DIS (tone onset at 0ms), for each DISdiff and MEMdiff condition (S1 and DIS), in the corresponding ROI represented on the MNI template. The strip plots show individual data averaged by condition in the gray periods showing positive or greater evidence (see Table 1 for time-windows and results). Lines connect data from the same participant. Only contralateral tones are analysed here: S1-R and DIS-R for left sources, and S1-L and DIS-L for right sources. S1-L or S1-R: S1 melodies presented to the left or right ear, respectively. DIS-L or DIS-R: DIS melodies presented to the left or right ear, respectively.

In the left Temporal ROI, the best model explaining N1 amplitude is the full model, including the triple interaction between StimulusType, MEMdiff, and DISdiff (BF_10_ = 222797.5). The analysis of effects reveals decisive evidence for StimulusType (BF_inclusion_ = 17470.0), with larger N1 for S1 compared to DIS, and strong evidence for the triple interaction (BF_inclusion_ = 13.4), which we further explore in separate analyses by Memory difficulty (for all other effects, BF_inclusion_<2.4). *Under low Memory difficulty*, in a Bayesian ANOVA with StimulusType and DISdiff as factors, the best model explaining N1 amplitude is the full model including the StimulusType by DISdiff interaction (BF_10_ = 118659.4). The analysis of effects reveals decisive evidence for StimulusType (BF_inclusion_ = 1692.7) and for the StimulusType by DISdiff interaction (BF_inclusion_ = 108.4), and weak evidence against DISdiff (BF_inclusion_ = 0.65). Bayesian t-tests exploring this interaction reveal strong evidence for an effect of DISdiff on N1 amplitude for S1 (BF_10_ = 51.3, larger N1 amplitudes for the easyDIS condition), but weak evidence for an absence of effect of DISdiff on N1 amplitude for DIS (BF_10_ = 0.36). *Under high Memory difficulty*, in a Bayesian ANOVA with StimulusType and DISdiff as factors, the best model explaining N1 amplitude is the model with only the StimulusType effect (BF_10_ = 104513.0), followed by the model with StimulusType and DISdiff effects (BF_10_ = 56277.9). The analysis of effects reveals decisive evidence for StimulusType (BF_inclusion_ = 103549.6) but weak evidence against DISdiff (BF_inclusion_ = 0.54).

In the right Temporal ROI, the best model explaining N1 amplitude is the model with StimulusType, DISdiff, and the interaction between StimulusType and DISdiff (BF_10_ = 4173.9), immediately followed by the model with the same effects and interaction and the effect of MEMdiff in addition (BF_10_ = 2399.1). The analysis of effects reveals decisive evidence for the effect of StimulusType (BF_inclusion_ = 1520.4), positive evidence for the interaction between StimulusType and DISdiff (BF_inclusion_ = 6.0), and weak evidence for an absence of an effect of MEMdiff (BF_inclusion_ = 0.69) and of DISdiff (BF_inclusion_ = 0.50). Further post-hoc Bayesian t-test to explore the StimulusType-by-DISdiff interaction reveal that, for S1, there was positive evidence for larger N1 amplitude for easyDIS than for hardDIS (BF_10_ = 4.2), whereas for DIS, there is weak evidence for no difference in N1 amplitude between easyDIS and hardDIS (BF_10_ = 0.63).

In the left Frontal ROI, the best model explaining N1 amplitude is the model with only the effect of StimulusType (BF_10_ = 5266.8). Other models have similar BF_10_, but the analysis of effects reveals only decisive evidence for StimulusType (BF_inclusion_ = 3088.3; for all other effects and interactions, BF_inclusion_ < 2.1).

In the right Frontal ROI, the best model explaining N1 amplitude is the model with StimulusType (BF_10_ = 22.3), followed by the model with StimulusType, DISdiff, and the interaction between StimulusType and DISdiff (BF_10_ = 17.1), the model with StimulusType and DISdiff (BF_10_ = 11.9), and the model with StimulusType, MEMdiff and the interaction between StimulusType and MEMdiff (BF_10_ = 8.4). The analysis of effects confirms strong evidence for the effect of StimulusType (BF_inclusion_ = 22.5), and only weak evidence for the interactions between StimulusType and DISdiff (BF_inclusion_ = 1.4) and between StimulusType and MEMdiff (BF_inclusion_ = 1.6); there is weak to positive evidence against all other effects (BF_inclusion_ < 0.55).

In the four ROIs, the N1 is larger for S1 than DIS, revealing efficient filtering. In the left temporal ROI, the N1 to S1 is larger in easyDIS than in hardDIS, under lowM only. In the right temporal ROI, the N1 to S1 is larger in easyDIS than in hardDIS, irrespective of the memory task difficulty.

#### Transient Evoked Responses: P2 to tones 2, 3, 4 (S1 and DIS)

Activity is observed in the contralateral Planum Polare in the P2 latency range (200-250 ms) for tone 2,3,4 (S1 and DIS). As for the N1, P2 amplitudes are analyzed with Bayesian repeated-measure ANOVAs with StimulusType, MEMdiff, and DISdiff as factors (see Table 1 and Figure 8).

**Figure 8.**
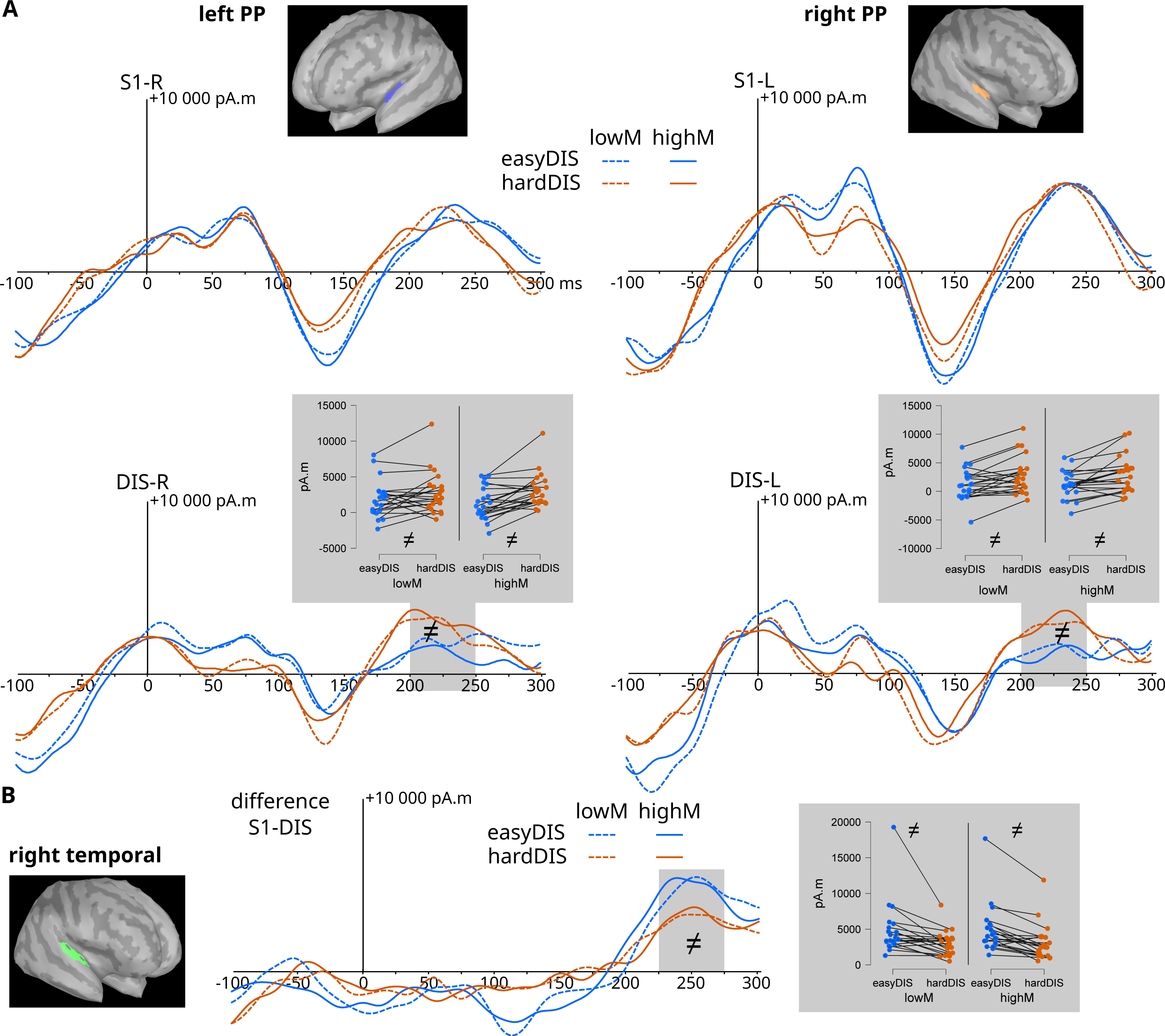
Transient evoked responses to the tones 2, 3 and 4 in the source space (P2 and late differential response ROIs). (A) Time course of the averaged ERF to the tones 2, 3 and 4 of the to-be-attended melody S1 and of the to-be-ignored melody DIS (tone onset at 0ms), for each DISdiff and MEMdiff condition (S1 and DIS), in the corresponding ROI represented on the MNI template. (B) Time course of the difference (S1-DIS) between averaged ERF to the tones 2, 3 and 4 of the to-be-attended melody S1 and of the to-be-ignored melody DIS (tone onset at 0ms), for each DISdiff and MEMdiff condition, in the corresponding ROI represented on the MNI template. The strip plots show individual data averaged by condition in the gray periods showing positive or greater evidence (see Table 1 for time-windows and results). Lines connect data from the same participant. Only contralateral tones are analyzed here: S1-R and DIS-R for left sources, and S1-L and DIS-L for right sources. S1-L or S1-R: S1 melodies presented to the left or right ear, respectively. DIS-L or DIS-R: DIS melodies presented to the left or right ear, respectively. leftPP: left Planum Polare, right PP: right Planum Polare.

In the left Planum Polare, the best model explaining P2 amplitude is the model with StimulusType, DISdiff, and the interaction between StimulusType and DISdiff (BF_10_ = 32.1). The analysis of effects confirms positive evidence for the interaction between StimulusType and DISdiff (BF_inclusion_ = 5.6) and for the effect of DISdiff (BF_inclusion_ = 5.0), and weak evidence for the effect of StimulusType (BF_inclusion_ = 1.1). Further post-hoc Bayesian t-test to explore the StimulusType-by-DISdiff interaction reveal that, for S1, there is positive evidence for no difference in P2 amplitude between easyDIS and hardDIS (BF_10_ = 0.23), whereas for DIS, there is strong evidence for a larger P2 amplitude in hardDIS compared to easyDIS (BF_10_ = 22.8).

In the right Planum Polare, the best model explaining P2 amplitude is the model with StimulusType, DISdiff, and the interaction between StimulusType and DISdiff (BF_10_ = 855.6), followed by the model with StimulusType and DISdiff (BF_10_ = 732.2). The analysis of effects confirms an effect of StimulusType with decisive evidence (BF_inclusion_ = 123.0; P2 is larger for S1 than DIS), of DISdiff with positive evidence (BF_inclusion_ = 6.9), and of the interaction between StimulusType and DISdiff with weak evidence (BF_inclusion_ = 1.2). Further post-hoc Bayesian t-test to explore the StimulusType-by-DISdiff interaction reveal that, for S1, there is positive evidence for no difference in P2 amplitude between easyDIS and hardDIS (BF_10_ = 0.25), whereas for DIS, there is strong evidence for a larger P2 amplitude in hardDIS compared to easyDIS (BF_10_ = 69.9).

33

#### Transient Evoked Responses: late differential response to tones 2, 3, 4 (S1 - DIS)

The ERF evoked by S1 and DIS not only differs during N1 and P2, as described above, but they also diverge later, between 225 and 275 ms, in right temporal areas around Heschl’s gyrus, irrespective of stimulation side. This late differential response (S1 – DIS amplitude) is examined with Bayesian repeated-measure ANOVAs with Stimulus_Side, DISdiff, and MEMdiff as factors (see Table 1 and Figure 8).

In the right Temporal ROI, the best model explaining the late differential response (S1 – DIS amplitude) is the model with the factors Stimulus_Side, DISdiff, and the interaction between Stimulus_Side and DISdiff (BF_10_ = 91.2), followed by the model with only the effect of DISdiff (BF_10_ = 75.5) and the model with the effects of Stimulus_Side and DISdiff (BF_10_ = 36.4). The analysis of effects confirms strong evidence for DISdiff (BF_inclusion_ = 42.9) and weak evidence for the interaction between Stimulus_Side and DISdiff (BF_inclusion_ = 2.0), while there is weak evidence against an effect of Stimulus_Side (BF_inclusion_ = 0.49). The difference between S1 and DIS ERF amplitude is larger for easyDIS than hardDIS.

#### Sustained Evoked Responses

During S1 and DIS presentation, irrespective of S1 presentation side, emergence tests (time window 500-2000 ms after S1 onset) reveal activity in bilateral temporal areas, in two distinct clusters in the Planum Polare and the Planum Temporale. Activity in these temporal ROIs is examined with Bayesian repeated-measure ANOVAs with S1side, DISdiff, and MEMdiff as factors (see Table 1 and Figure 9).

**Figure 9.**
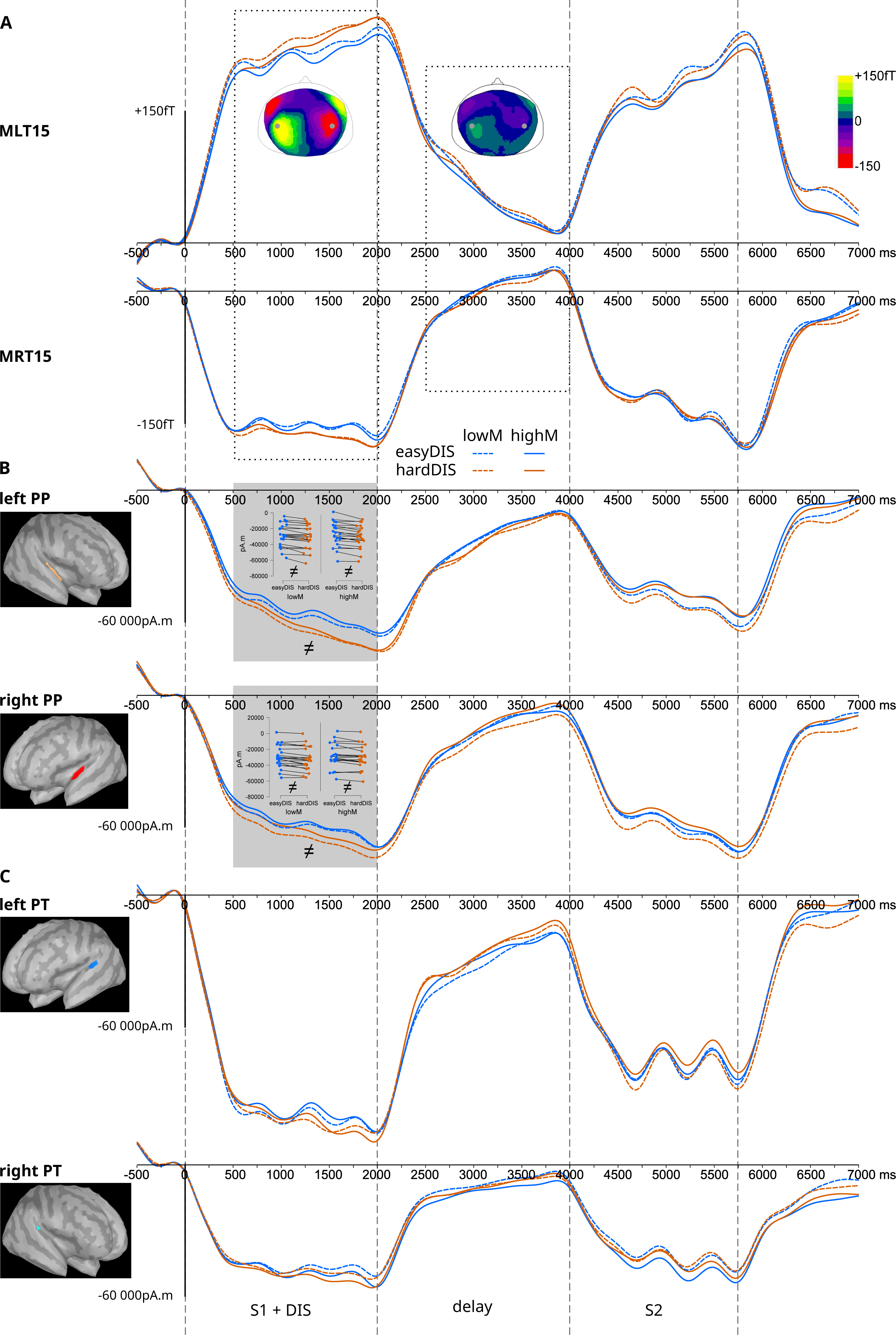
Sustained evoked responses. (A) In the sensor space, time course of the ERF to S1 & DIS melodies, the delay and to S2 (S1 onset at 0ms), for each DISdiff and MEMdiff condition, at MLT15 and MRT15 (above the left and right temporal lobes, respectively). Topoplots show sensor data over the time periods of interest, indicated by the dotted rectangles (500 to 2000 during S1 & DIS melodies, 2500 to 4000 during the delay). Gray dots on topographies indicate corresponding sensor position. (B & C) In the source space, time course of the ERF for each DISdiff and each MEMdiff condition, in the corresponding ROIs represented on the MNI template. The strip plot shows individual data averaged by condition in the gray periods showing positive or greater evidence (see Table 1 for time-windows and results). Lines connect data from the same participant.

**Figure 10.**
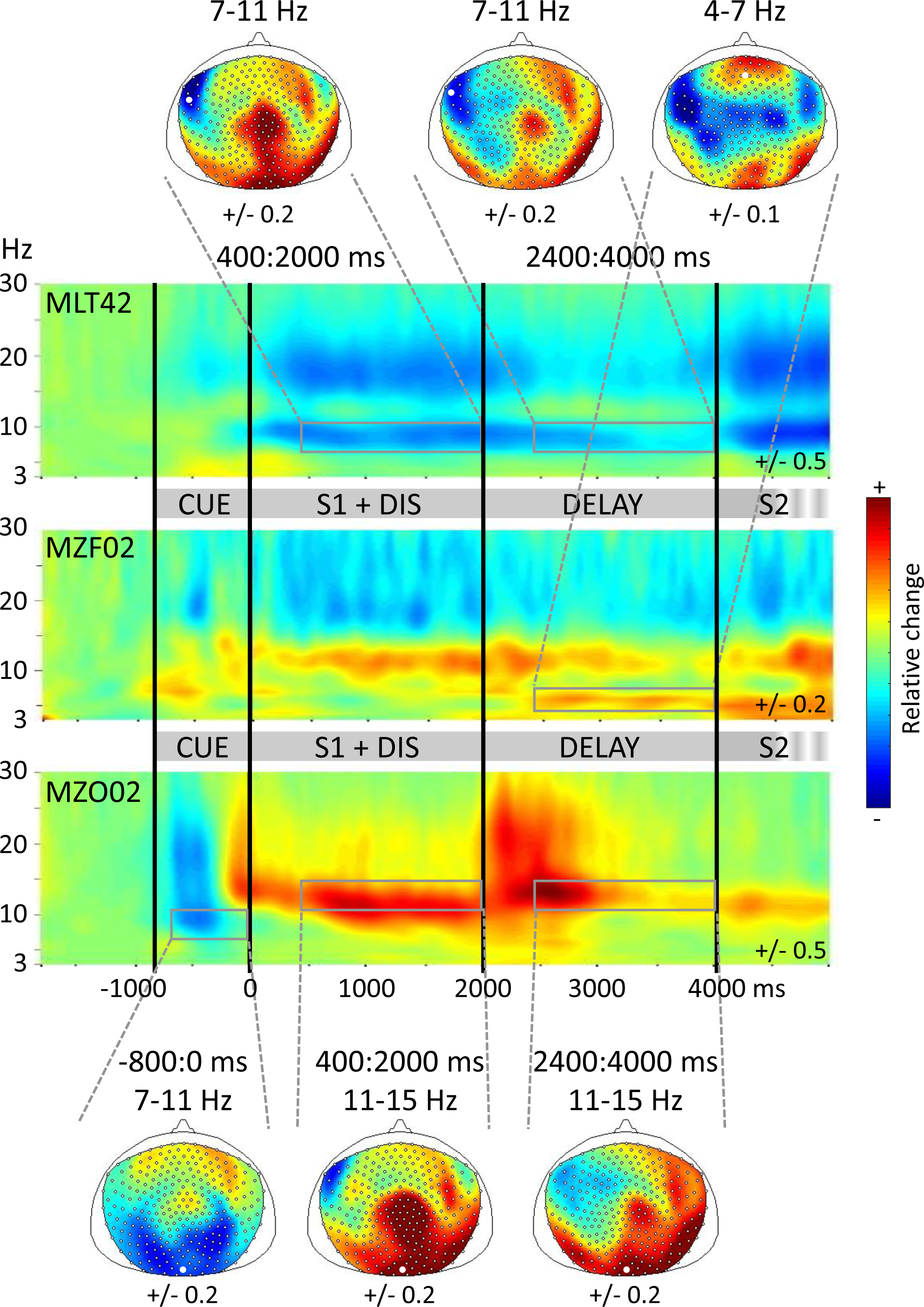
Time-frequency plots in the sensor space. Time-frequency power (relative change) plots during Cue presentation, S1 & DIS melodies, and the delay (S1 onset at 0ms), for all conditions averaged, at a temporal (MLT42), a frontal (MZF02) and an occipital sensor (MZO02) and corresponding topographies. Response to S2 is not fully represented here. In comparison to baseline (1800 to -1000 ms relative to S1 onset), a decrease in power appears in blue and an increase in power appears in red. White dots on topographies indicate corresponding sensor position.

In the left Planum Polare, the best model explaining results during presentation of S1 and DIS melodies is the one with the factors DISdiff and S1side (BF_10_ = 129061.0), immediately followed by the model with only DISdiff (BF_10_ = 94853.0), and several other models with similar evidence. The analysis of effects only reveals decisive evidence for the effect of DISdiff (BF_inclusion_ = 89087.1) and weak evidence for the effect of S1side (BF_inclusion_ = 1.4), whereas there is weak to positive evidence against all other effects (BF_inclusion_ < 0.63). Sustained evoked responses in the left Planum Polare during S1 and DIS are larger in condition hardDIS compared to easyDIS.

In the right Planum Polare, the best model explaining sustained evoked response amplitudes during S1 and DIS is the one with only the factor DISdiff (BF_10_ = 207.1). Other models explain almost as well the data, however the analysis of effects brings only decisive evidence for an effect of DISdiff in the right Planum Polare (BF_inclusion_ = 202.2). As in the left hemisphere, response amplitudes are larger in condition hardDIS compared to easyDIS.

In the left Planum Temporale, the best model explaining sustained evoked response amplitude during S1 and DIS is the one with factors DISdiff, MEMdiff, S1side, and the interaction between DISdiff and MEMdiff (BF_10_ = 1.6). Several other models, including the null model, explain almost as well the data. The analysis of effects only reveals weak evidence for the effect of DISdiff (BF_inclusion_ = 1.9) and S1side (BF_inclusion_ = 2.2), with differences in the same direction as in the Planum Polare, and weak evidence for no effect of MEMdiff (BF_inclusion_ = 0.90) and of the interaction between DISdiff and MEMdiff (BF_inclusion_ = 0.47).

In the right Planum Temporale, the best model explaining sustained evoked response amplitude during S1 and DIS is the null model (for all other models, BF_10_ < 0.65).

During the retention delay (time window 2500-4000 ms), sustained evoked responses are observed only in the left hemisphere, irrespective of S1 side, in the same areas as during S1 and DIS presentation. Activity in these left temporal ROIs is examined with Bayesian repeated-measure ANOVAs with S1side, DISdiff, and MEMdiff as factors.

In the left Planum Polare, the best model explaining sustained response amplitude is the null model (for all other models, BF_10_ < 0.57).

In the left Planum Temporale, the best model explaining sustained response amplitude is the model with only the S1side effect (BF_10_ = 1.13). Other models, including the null model, explain almost as well the data, and the analysis of effects reveals only weak evidence for the effect of S1side (BF_inclusion_ = 1.13), with larger amplitudes ipsilaterally.

### Time-Frequency results

#### Sensor-level activation

The time-frequency power (relative change) plots computed on sensor-level data (see **Error! Reference source not found.**) with a baseline from -1.8 to -1s relative to S1 onset reveal a decrease in alpha power after cue onset at occipital sensors. A decrease in alpha power (7-11 Hz) is observed during the presentation of the S1 and DIS melodies and during the delay at left temporal sensors; while an increase in alpha power (11-15 Hz) is observed at occipital sensors. These two alpha sub-bands will be named low and high alpha in the following. Finally, an increase in theta power (4-7 Hz) is observed during the retention delay at frontal sensors.

#### Source-level whole brain activation

Sources of theta and alpha activities are estimated using DICS beamformer in different time-frequency windows of interest, and contrasted to a pre-cue baseline period using non-parametric cluster-based permutation testing, for S1 presented to the left or right ear separately (Figure 11).

**Figure 11.**
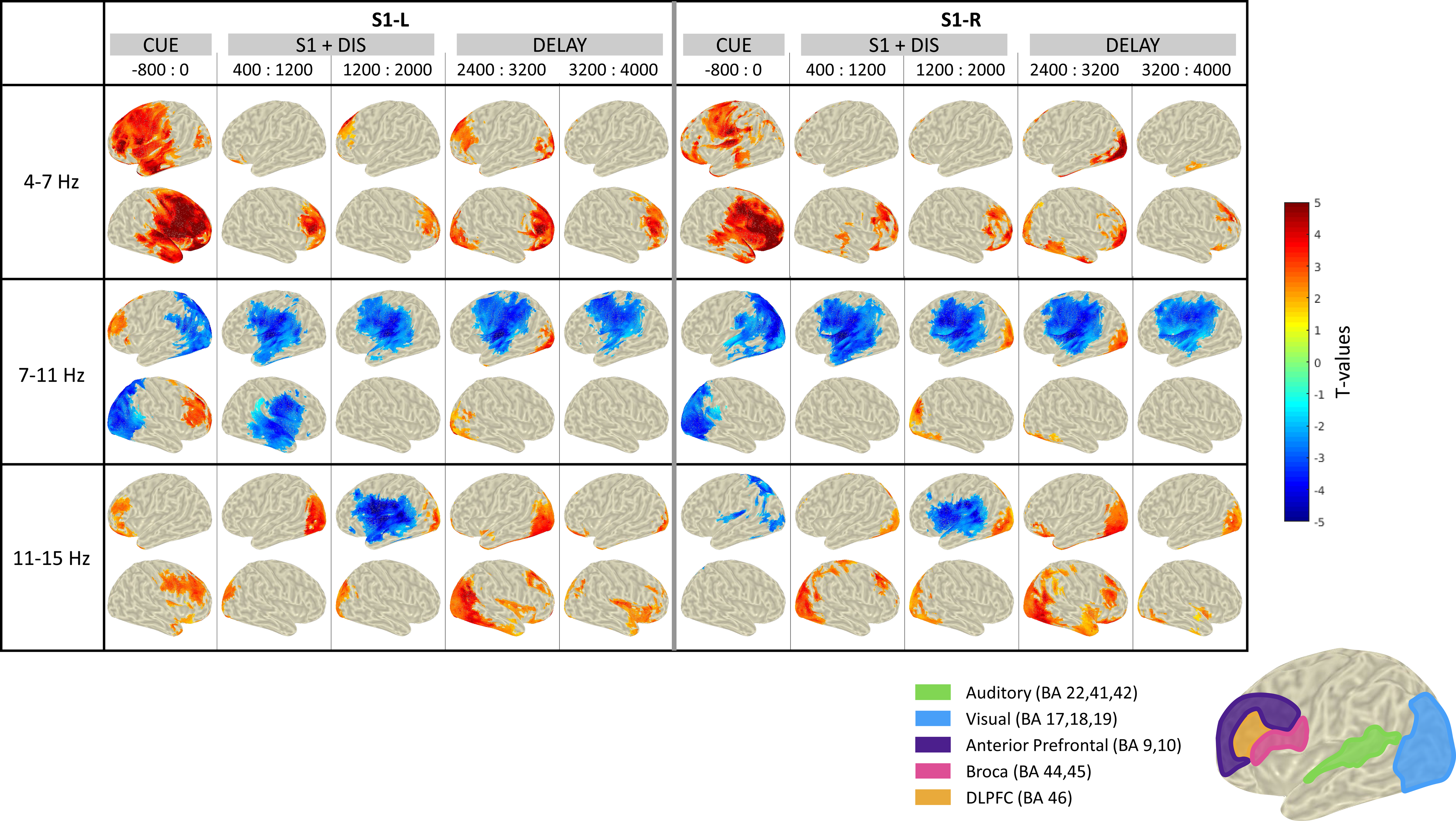
Source activity in theta and alpha bands. Distributions of T values for each time-frequency window in comparison to a pre-cue baseline (-1800ms to -1000ms locked to S1 onset) for trials with S1 presented to the left or the right, separately (S1-L & S1-R). The results are masked with the p-value coming from the cluster-based permutation tests: only significant clusters are shown (p<0.05). In comparison to baseline, a decrease in power appears in blue and an increase in power appears in red. In the right bottom corner, the chosen regions of interest for further analyses are presented. The corresponding Brodmann areas are indicated in brackets for each region of interest.

The main power modulations are:

1. An increase in theta power in anterior prefrontal regions (BA 9, 10) during cue, melody presentation, and delay, more pronounced in the right hemisphere (Figure 11, top row);
2. A decrease in low alpha power in auditory regions in superior temporal cortices (BA 22, 41, 42), Broca area (BA 44, 45), and dorsolateral prefrontal cortex (DLPFC, BA 46) during melody presentation in both hemispheres, and during the delay in the left hemisphere (Figure 11, middle row);
3. A decrease in low alpha power in visual regions in occipital cortices (BA 17, 18, 19) during cue presentation (Figure 11, middle row) followed by an increase in high alpha power during melody presentation and delay (Figure 11, bottom row).

#### Source-level ROI activation

According to the analysis of the individual peak frequency in each ROI, we investigated the following frequency bands: 4-7 Hz in anterior prefrontal regions; 8-11 Hz for decrease in alpha power in the auditory, visual, Broca and DLPFC ROIs; and 9-12 Hz for increase in alpha power in the visual ROI. To investigate the impact of the memory task and distractor difficulties, power values in time-frequency windows of interest are examined using Bayesian repeated-measure ANOVAs with S1side, DISdiff, and MEMdiff as factors (see Table 2 and Figure 12).

**Figure 12.**
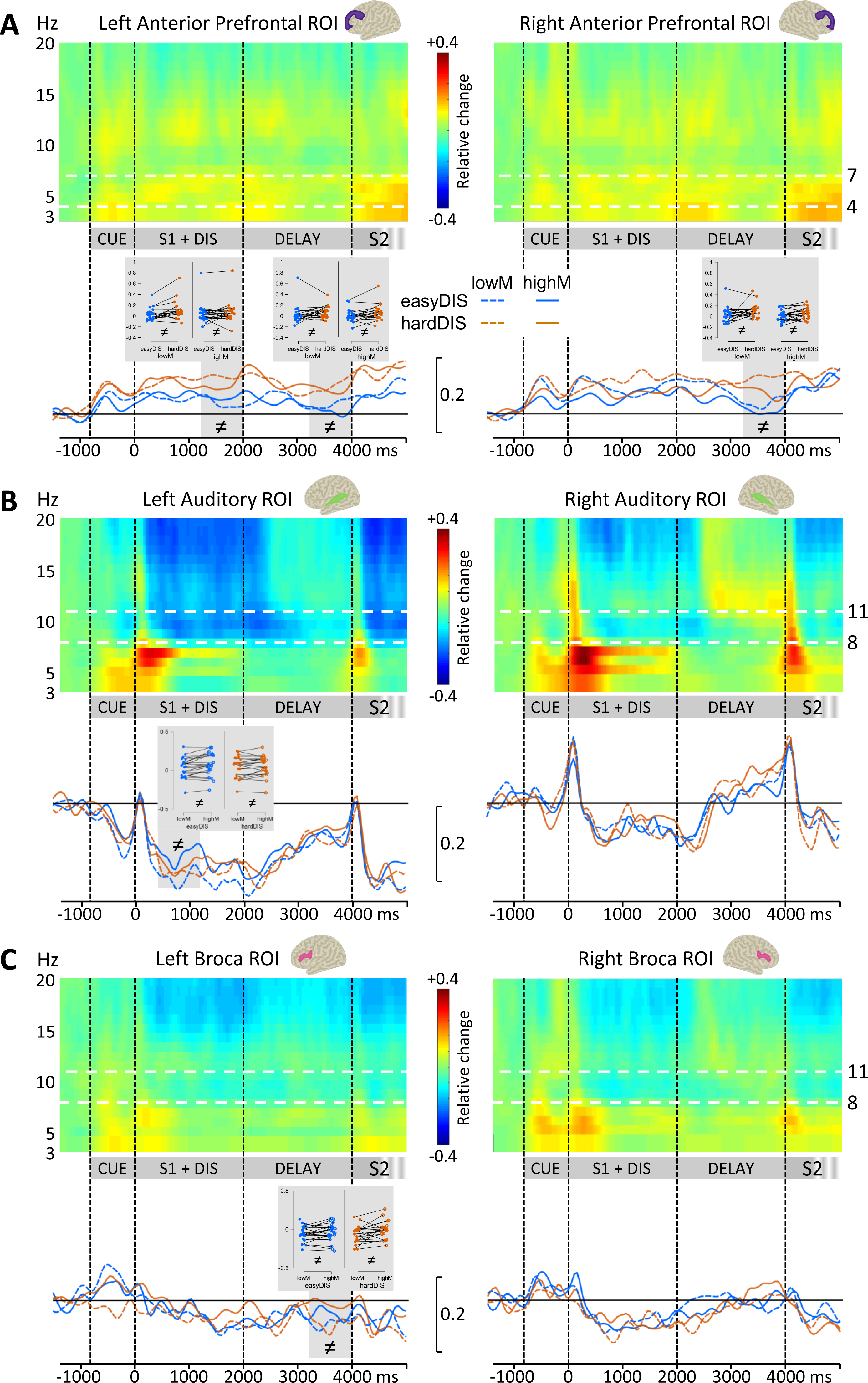
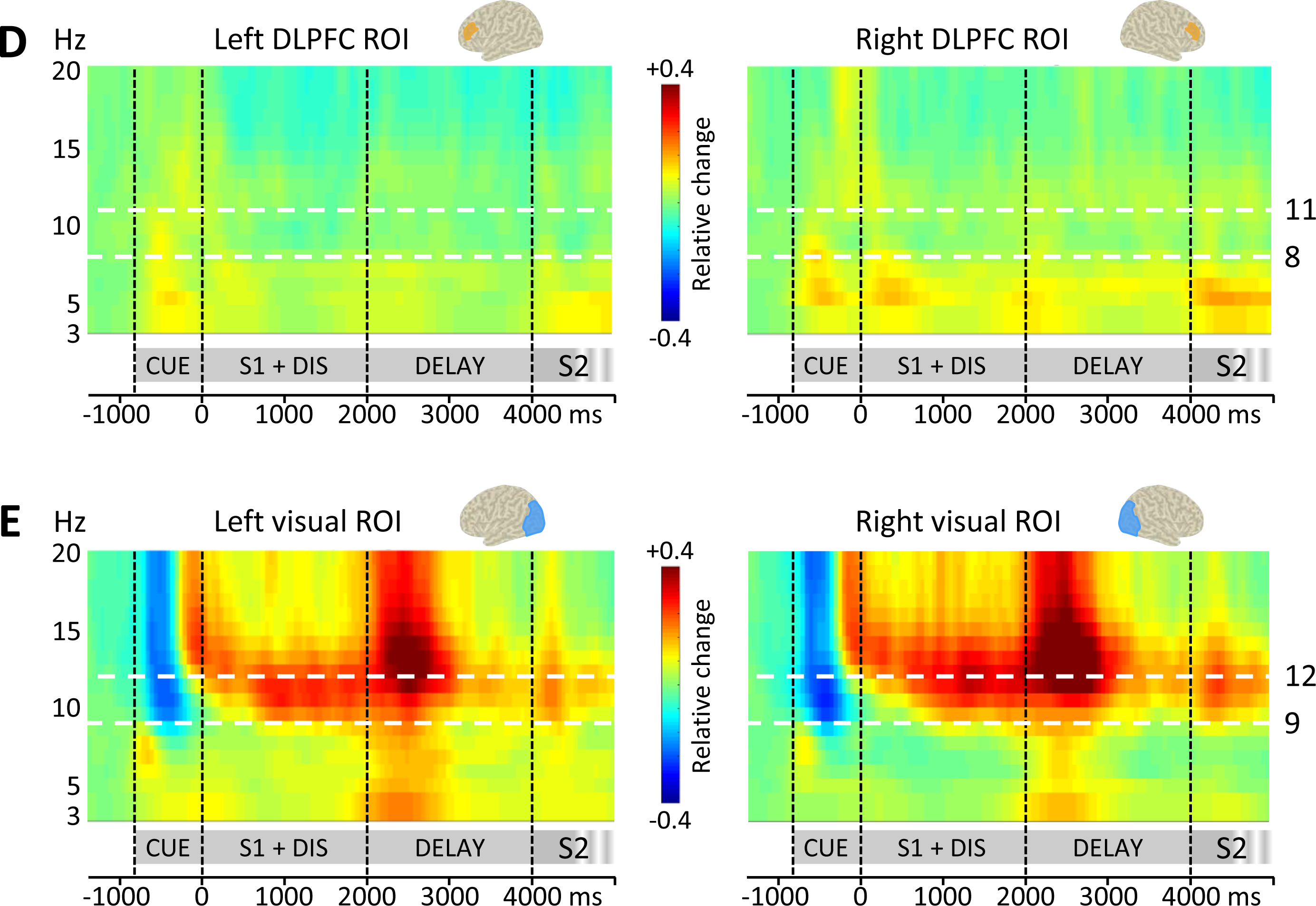
Time-frequency plots and time profiles in the source space. Time-frequency power (relative change) plots during Cue presentation, S1 & DIS melodies, and the delay (S1 onset at 0ms), for all conditions averaged, in the left and right Anterior Prefrontal regions (A), in the left and right Auditory cortices (B), in the left and right Broca areas (C), in the left and right Dorso-Lateral Prefrontal Cortex (DLPFC, D), and in the left and right Visual cortices (E). Response to S2 is not fully represented here. In comparison to baseline (-1800 to -1000 ms relative to S1 onset), a decrease in power appears in blue and an increase in power appears in red. Regions of interest were defined based on the results from the cluster-based permutation tests performed on the DICS-reconstructed sources for the whole brain. Time course of the power (relative change) for each DISdiff and each MEMdiff condition, in the corresponding ROIs represented on the MNI template: alpha band (8 to 11 Hz) in the Auditory (A) and BROCA (B) ROIs, theta band (4 to 7 Hz) in the Anterior Prefrontal (C) ROIs. The strip plot shows individual data averaged by condition in the gray periods showing positive or greater evidence (see Table 2 for time-windows and results). Lines connect data from the same participant.

*Anterior prefrontal theta*. In the anterior prefrontal regions, the best model explaining results in the 4-7 Hz band after cue onset and before S1 onset is the one with the factors DISdiff and S1side in the left hemisphere (BF_10_ = 5.0) and with the factor MEMdiff in the right hemisphere (BF_10_ = 1.2). The analysis of effects, in the left hemisphere, revealed positive evidence for the effect of S1side (BF_inclusion_ = 3.7) and weak evidence for the effect of DISdiff (BF_inclusion_ = 1.4). The increase in theta power is larger after the left cue presentation. The best model explaining results in the 4-7 Hz band after S1 onset is the one with the factor DISdiff in the left hemisphere (BF_10_ = 1.9 between 400 and 1200 ms; BF_10_ = 3.3 between 1200 and 2000 ms; BF_10_ = 4.5 between 3200 and 4000 ms) and in the right hemisphere (BF_10_ = 10.6 between 3200 and 4000 ms); otherwise the null model is the best model (for all other models, BF_10_<0.89). The increase in theta power is larger in the hardDIS condition.

*Auditory and prefrontal low alpha.* In the left auditory regions, the best model explaining results in the 8-11 Hz band during the presentation of S1 and DIS melodies is the one with the factor MEMdiff only (BF_10_ = 3.4 between 400 and 1200 ms, BF_10_ = 1.7 between 1200 and 2000 ms), and during the delay, it is the null model. The decrease in alpha power is larger in the lowM condition.

In the right auditory regions, the best model explaining results in the 8-11 Hz band during presentation of S1 and DIS melodies is the one with the factor S1side (BF_10_ = 1.5 between 400 and 1200 ms). Between 1200 and 2000ms, the best model is the null model (for all other models, BF_10_<0.94).

In the Broca areas, the best model explaining results in the 8-11 Hz band after S1 onset is the null model (for all other models, BF_10_<0.41), except between 400 and 1200 ms (best model with factor S1side: BF_10_ = 1.1) and between 3200 and 4000 ms (best model with factor MEMdiff: BF_10_ = 3.2) in the left hemisphere. The decrease in alpha power is larger in the lowM condition.

In the left DLPFC, the best model explaining results in the 8-11 Hz band after S1 onset is the null model (for all other models, BF_10_<0.54).

*Visual low and high alpha.* In the bilateral visual regions, the best model explaining results in the 8-11 Hz band during cue presentation is the null model. In the bilateral visual regions, the best model explaining results in the 9-12 Hz band during S1 and DIS melodies and during the delay is the null model, except in the right visual regions between 400 and 1200 ms (best model with factor S1side: BF_10_ = 1.0) and between 3200 and 4000 ms (best model with factors DISdiff and S1side: BF_10_ = 2.6) (for all other models, BF_10_<0.77). The decrease after the cue and the increase after S1 onset in alpha power do not seem to be modulated by any of the investigated factors.

## Discussion

In the present study, we investigate auditory working memory and selective attention, in a delayed-matching-to-sample task (DMST) requiring to filter out distractors during encoding of melodic sequences in memory. Behavioral and MEG results allow to further understand working memory, auditory selective attention, and their interactions, as discussed below.

### Electrophysiological markers of non-verbal auditory working memory

During the DMST at the core of the MEMAT paradigm, we observe a series of electrophysiological markers that were expected based on prior studies: (1) following the presentation of the visual cue (which itself generated visual ERFs and an alpha decrease in visual areas), in anticipation of the sequence to encode, a slow magnetic field develops bilaterally (magnetic counterpart of the CNV, see below), notably in the Planum Polare, and theta power increases in anterior prefrontal regions; (2) transient and sustained evoked responses are observed during sequence encoding in superior temporal and inferior frontal areas, along with an increase in theta power in anterior prefrontal areas, a decrease in alpha power in a fronto-temporal network, and an increase in alpha power in visual areas; (3) a sustained evoked response persists in superior temporal areas during the retention delay, also along with an increase in theta power in anterior prefrontal areas, a decrease in alpha power in a left fronto-temporal network, and an increase in alpha power in visual areas.

This pattern of transient and sustained ERFs is consistent with previous descriptions in auditory non-verbal DMST (Albouy et al., 2013; Grimault et al., 2014). Here the addition of a visual cue at a fixed delay prior to S1 onset allows to characterize the slow response developing in anticipation of the sound sequences, that could reflect the sensory contribution to the CNV (see Brunia & Van Boxtel, 2001) and strongly resembles the auditory preparatory potential observed with intracranial recordings in epileptic patients slightly more posteriorly in Heschl gyri (Hu et al., 2020).

Power in the alpha band is modulated along the course of a trial: a transient decrease in visual areas following the visual cue, and then a sustained increase in visual areas associated with a sustained decrease in fronto-temporal areas during encoding and maintenance (we did not study the retrieval phase). Following the interpretation of the alpha rhythm as an inhibitory process (e.g., Klimesch et al., 2007; Foxe & Snyder, 2011; Jensen & Mazaheri, 2010; Thut et al., 2006; Palva & Palva, 2011), this pattern suggests a transient activation of the visual system to process the visual cue, followed by an inhibition of task-irrelevant occipital visual areas and an activation of the task-relevant fronto-temporal auditory network during sound encoding and maintenance in memory. In keeping with ElShafei et al. (2018, 2020), alpha oscillations are centered around a higher frequency in visual areas compared to auditory areas during auditory processing.

Power in the theta band increases in anterior frontal areas during cue processing and the subsequent encoding and maintenance phases. Studies of short-term and working verbal memory with EEG/MEG have almost always uncovered increased frontal theta oscillations, in particular during the maintenance phase (review in Pavlov & Kotchoubey, 2022). As discussed in Hsieh and Ranganath (2014), the specific role of theta oscillations (and theta-gamma coupling) in short-term/working memory processes is debated, and might further depend on the regions involved (frontal vs. hippocampal areas for example). Midline frontal theta oscillations have been in particular associated with maintaining temporal relationships between items (Hsieh et al., 2011). The present study reveals that frontal theta oscillations are also involved in non-verbal auditory memory, both during encoding and maintenance of pitch sequences.

Here, we manipulate memory task difficulty at the block level, and participants were warned about the difficulty of the upcoming block of trials. At the behavioral level, as expected (Blain, Talamini, et al., 2022; Blain, de la Chapelle, et al., 2022), accuracy decreases and response time increases when the difficulty of the memory task increases. At the neurophysiological level, an increase in the amplitude of the slow ERF (CNV) in anticipation of the target melody is observed in the high difficulty memory task relative to the low difficulty one when the left ear is cued, in the contralateral right auditory cortex (Figure 4). This result confirms the efficiency of the manipulation of task difficulty, with more pronounced anticipation of the relevant sound when the memory task is more difficult.

Beyond this anticipation effect, a smaller decrease in alpha oscillations was observed in the high difficulty memory condition relative to the low difficulty one, in the left auditory cortex during encoding of the S1 melody in memory (Figure 12B), and in the left inferior frontal cortex (Broca’s area) during the retention delay (Figure 12C). These results in the alpha band suggest that in the difficult memory condition, the left auditory fronto-temporal network is less recruited than during the easier memory condition. We will return to this point when discussing the interaction between memory and attention.

### Deployment of auditory selective attention during a working memory task

During sequence encoding in MEMAT, a distracting melody is presented in the other ear, which is either easy or hard to ignore (on different trials within the same blocks). Participants succeed in selecting to-be-attended (S1) over to-be-ignored (DIS) melodies as evidenced by larger ERFs for target tones over distracting tones in temporal and frontal regions (N1, P2, late response for notes 2-3-4, see Figures 5, 6, 7, 8 and Table 1; review in Giard et al., 2000). The difficulty of the attentional filtering impedes as expected on performance and reaction times (Blain, Talamini, et al., 2022; Blain, de la Chapelle, et al., 2022), with poorer performance and longer RTs for difficult-to-ignore distractors. This filtering difficulty modulates several of the electrophysiological markers listed above (see Tables 1 and 2), during the encoding phase of the DMST (transient and sustained ERFs, theta oscillations), as well as during the maintenance phase (theta oscillations). These modulations of electrophysiological responses shed light on how selective attention operates during a working memory task.

In response to the first distracting tone, a larger N1 amplitude is observed in response to the hard-to-ignore distractor than to the easy-to-ignore distractor in the left temporal areas (Figure 5). As in MEMAT we manipulate the frequency similarity between to-be-attended and to-be-ignored sounds to create two levels of filtering difficulty, the hard-to-ignore sounds being closer in frequency to the to-be-attended melody could catch more attention and be more difficult to suppress than the easy-to-ignore sounds, in line with contingent stimulus-driven orienting (Folk et al., 1992; Corbetta & Shulman, 2002). This suggests a more efficient inhibition of the easy-to-ignore distracting tones as observed in previous EEG studies of auditory selective attention (Alain & Woods, 1994; Melara et al., 2005).

Critically, in bilateral superior temporal regions, in response to subsequent tones, N1 amplitude for the to-be-attended tones is larger when accompanied with easy-to-ignore distracting melodies, whereas the N1 to the to-be-ignored tones is not modulated by filtering difficulty (Figure 7). It suggests a facilitation of the processing of the target, more efficiently so when the distracting sounds are dissimilar to the target, in line with previous findings (Alain & Woods, 1994; Alain et al., 1993).

Later on, more anteriorly, in bilateral planum polare, the amplitude of the P2 to the tones of the melody to ignore is smaller when these tones were easy rather than difficult to ignore, in keeping with previous results (Alain & Woods, 1994). This suggests a more efficient inhibition of the distracting tone processing when they are easy to ignore (Figure 8).

Around 250 ms, a larger ERF amplitude for to-be-attended tones than for to-be-ignored tones in right temporal areas seems to be related to an additional component superimposed to the obligatory P2, such as the PN (« Processing Negativity »; Näätänen, 1982, 1992). This effect is more pronounced when the to-be-ignored tones are easy to ignore, i.e. more dissimilar to the to-be-attended tones (Alho et al., 1987) or when the task is easier (Melara et al., 2005). This increased component suggests a larger differential processing of relevant and irrelevant tones when distractors are easy to ignore, and has been interpreted as reflecting active suppression of irrelevant signal (Alho et al., 1987; Melara et al., 2005).

Taken together, these findings highlight that distinct attentional mechanisms are at play, with facilitation of target processing preceding inhibition of irrelevant tones, in keeping with previous findings (Michie et al., 1993; Alho et al., 1987, 1994; Bidet-Caulet et al., 2007, 2010; Alain & Woods, 1994; Alain et al., 1993; Melara et al., 2005). These two effects have distinct localizations: auditory areas in the vicinity of the primary auditory cortex for target facilitation, and more anterior superior temporal areas in the planum polare for distractor inhibition, in line with previous observations (Rif et al., 1991). Except for the later differential effect (∼250 ms) which was observed only in the right hemisphere, all these effects on transient ERFs were bilateral in agreement with most previous studies. All these mechanisms are less efficient when targets and distractors are more similar (Alain & Woods, 1994; Alain et al., 1993).

During sound sequence encoding, in addition to transient ERFs following each tone of the target or distracting melodies, a sustained evoked response was observed in bilateral superior temporal areas. In bilateral planum polare, the sustained evoked response is larger when the distractors are hard-to-ignore compared to easy-to-ignore, suggesting that this component reflects the deployment of cognitive resources (listening effort) to solve the difficult filtering task. A similar modulation of the sustained evoked response during S1 encoding by task difficulty in an auditory DMST was also observed in Albouy et al. (2013). It is noteworthy here that sustained evoked responses and target-related transient ERFs are modulated in opposite directions by the difficulty to filter out the distracting melodies, suggesting that the deployment of additional resources when the distractors are hard-to-ignore (reflected in the sustained response) is not sufficient to restore the same level of differential processing between targets and distractors in auditory areas (reflected in target and distractor-related transient ERFs). This failure translates in behavioral data as costs of filtering difficulty, both for d’ and RT.

These effects on the ERFs highlight that during sound sequence encoding in memory, the processing of targets (and distractors) is modulated by attentional processes, possibly resulting in more or less precise memory traces of target sounds, which could impact on performance in the retrieval phase. Furthermore, we observe that theta power during sound sequence encoding and retention is larger when the distracting melodies is hard to ignore (compared to when they were easy to ignore). As described above, frontal theta oscillations have been associated with working memory processes including the processing of serial order in sequences (Hsieh et al., 2011). The current result would thus imply that a stronger involvement of these time-based memory processes is necessary when parsing interleaved target and distracting sounds is more demanding. This increase in theta power with increasing filtering difficulty is also in line with a link between theta activity and increased effort (Wisniewski et al., 2017).

### Interactions between auditory selective attention and working memory

At the behavioral level, we observe, as in Blain, de la Chapelle et al. (2022) and Blain, Talamini et al. (2022), an interaction between filtering difficulty and the memory task difficulty on performance (d’), with greater differences between easy and hard-to-ignore distractors in the low difficulty memory task than in the high difficulty memory task. In Blain, Talamini, et al. (2022), based on several lines of evidence derived from behavioral measures in non-musician and musician participants, we have argued that this interaction observed in d’ reflects shared cognitive resources between auditory selective attention and working memory, compatible with the *cognitive load theory* (Lavie, 2005; Lavie et al., 2004; Yi et al., 2004).

In the neurophysiological data, we observe an interaction between filtering difficulty and the memory task difficulty on the amplitude of the N1 to to-be-attended sounds in the left auditory cortex during sequence encoding, with a difference of target N1 amplitude according to filtering difficulty only in the low difficulty memory task. This neurophysiological result exactly mirrors the behavioral data (Figures 3 and 7). It suggests that under low memory load, attentional filtering is more efficient, resulting in larger response amplitudes for the to-be-attended sounds when the distractors are easy to ignore. Conversely, under high memory load, less resources would be available for attentional filtering, resulting in a reduced facilitation of relevant sound processing when the distractors are easy to ignore. Interestingly this interaction between filtering difficulty and memory task difficulty on the amplitude of the N1 to target sounds is only observed in the left auditory cortex, where a smaller decrease in alpha power is observed under high memory task difficulty. Hence, in the left auditory areas, under high memory load, the level of excitability is lower (larger alpha power) and the attentional facilitation of transient evoked responses to relevant sounds is reduced when distracting sounds are hard-to-ignore. A possible interpretation is that the left auditory cortex plays a critical role in attentional filtering (Bidet-Caulet et al., 2007), and that increasing the cognitive load with enhanced memory task difficulty impedes on these processes. In the framework of the cognitive load theory, we could interpret this finding as attributing less cognitive resources to a process subserving selective attention.

Further evidence for an interplay between attentional filtering and memory processes was obtained by analyzing frontal theta oscillations. During both sound sequence encoding and the maintenance delay, the power of theta oscillations which are thought to reflect time-based serial order processes in memory tasks (Hsieh et al., 2011) was increased when target selection was difficult. This suggest that time-based processes, which are relevant both for information selection and retention could be shared between attention and memory.

### Conclusion

Here by manipulating the difficulties of an auditory memory task (DMST) and of attentional filtering of distracting sounds, we could shed light on the neurophysiological processes subserving working memory, selective attention, and their interactions in the auditory modality. Attentional selection during sound sequence encoding operates through a combination of target processing facilitation and distractor inhibition that are reduced with increasing filtering difficulty. Increasing memory demands results in better anticipation of the sound sequences to encode (higher CNV amplitude) but also in a smaller desynchronization of alpha oscillations in the left auditory and frontal areas which could hinder attentional filtering, as reflected in target-related ERFs. Conversely, increasing attentional demands increases sustained evoked activity in superior temporal areas, which could favor filtering, but also enhances theta oscillations in anterior frontal areas which could reflect the involvement of a general time-based processing system. Future work is needed to understand how the interplay between these different processes is orchestrated.

## Acknowledgments

We would like to thank Lesly Fornoni and Anne Cheylus for their technical support.

